# Differential expression analysis in single cell and spatial RNASeq without model assumptions

**DOI:** 10.1101/2025.10.20.683496

**Authors:** Gennady Margolin, Andrew Tang, Sergey Leikin

## Abstract

Gene up(down)regulation findings in single cell and spatial RNASeq can be inconsistent despite remarkable progress in technology. False findings in high-quality samples raise concerns about assumptions behind widely accepted data analysis approaches. We therefore propose a weighted averaging approach for data analysis without assuming anything besides randomness of technical noise. This approach is closely related to prior work on statistics of cluster-randomized experiments. We show that weighing transcript counts based on measured noise variances and utilizing weighted rather than standard unweighted tests reduces both false positive and false negative findings. Our approach eliminates the need for parametrizing data distributions and/or rescaling transcript counts, which may cause artifacts by distorting and biasing the data. The resulting analysis is less complex and produces more consistent differential gene expression estimates.

## INTRODUCTION

Identification of up– and downregulated genes from single cell (scRNASeq) and spatial (spRNASeq) transcriptomics is a key element of many biological studies. Statistical analysis of this differential gene expression (DGE) is increasingly important, yet false findings is a known problem in scRNASeq ^1,2^. Many different approaches to data analysis designed to increase statistical power while reducing false findings have been proposed. However, all commonly used ^3–7^ and more recent models and methods ^8–27^ for this analysis still rely on unnecessary simplifications, data fitting, and/or parametrization that may cause false findings. For instance, the Wilcoxon test does not take into account up to 100-fold variation in the number of detected transcripts from cell to cell. A commonly used “pseudo-bulk” approach involves an assumption that cells contribute proportionally to the number of transcripts detected in them, like in bulk RNASeq. We show that such implicit assumptions in common DGE analysis approaches are not consistent with experimental data. In scRNASeq, each cell is an independent experimental cluster, the number of detected transcripts is the cluster size, and effects of the cluster size and its variability have been well established before in statistics of cluster-randomized experiments ^28–31^.

We also find that parametrization of transcript distribution (e.g., with negative binomial one to rescale the counts in SCTransform ^7^) is not beneficial for the analysis of good quality scRNASeq or spRNASeq data. In silico modeling of the ever-present sequencing depth variation and other tests show that altering the original data based on parametrizations and widely used logarithmic normalization may produce significant artifacts in DGE analysis.

Here we propose a new approach to DGE analysis in sc– and sp-RNASeq, which makes no assumptions beyond random technical noise and sampling to more accurately describe the data. We use known features of scRNASeq physics to derive the needed data distribution moments directly from measured transcript counts without assuming a specific form of the data distribution or modeling it. We do not assume the same transcript counting accuracy in different cells, because the observed cell-to-cell variations in the total counts suggest highly variable accuracy. Instead, we deduce relative contributions of different cells to the mean gene expression and its variance directly from the data by utilizing well-established methods of unbiased weighted averaging ^32^.

Comparison of the results produced by our method with commonly used approaches revealed significant differences in the results. We therefore benchmarked all methods with simulated data, experimental data modified by well-defined changes in expression, and simpler count subsampling in experimental data that mimics variations in the sequencing depth. Based on the known ground truth, this statistical testing revealed many more false positive and false negative findings in the commonly used approaches compared to our one. We invite interested readers to examine our findings and judge the reliability of the output produced by the commonly used methods on their own. The annotated R code for our tests and simulations is available at https://github.com/sergeyleikin/sc-sp-RNASeq.

Understanding the basic principles behind our approach and its practical usage do not require any special expertise. We define all concepts and explain their meaning in the main text, while focusing on theoretical proofs and derivations in supplemental material. We implemented all procedures for our data analysis as a simple set of functions compatible with data objects generated by the popular Seurat package for R programming language (https://satijalab.org/seurat/). Benchmarking of their computational performance vs. commonly used Seurat functions is provided at the end of Supplement 1. Our set of functions and their source codes can be downloaded at https://github.com/sergeyleikin/sc-sp-RNASeq. For simplicity and consistency, we compare the results produced by our method with the standard models within the Seurat workflow, using publicly available 10X Genomics scRNASeq datasets (https://www.10xgenomics.com/datasets) and our spRNASeq data collected with the 10X Genomics Visium HD assay. We first formulate our arguments and tools for scRNASeq and then adapt them to spRNASeq at the end of the manuscript.

## RESULTS

### Statistical weights of cells significantly affect average expression of many genes

Single cell RNA sequencing (scRNASeq) data is a matrix *N_Z,i_* of unique gene *Z* transcript counts in cell *i* (UMI counts, see Supplement 1). Even functionally similar cells exhibit large (up to ∼ 100-fold) variation in the total number of counts per cell *N_i_* as illustrated in Fig. 1A for GABAergic and glutamatergic neurons from embryonic mouse brain. This variation may be affected by biological differences between cells and technical variability of the assay. Therefore, relative gene expression

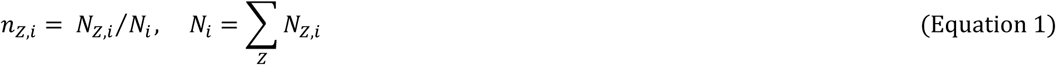

is recalculated from the data and further used in all statistical models.

**Figure 1.**
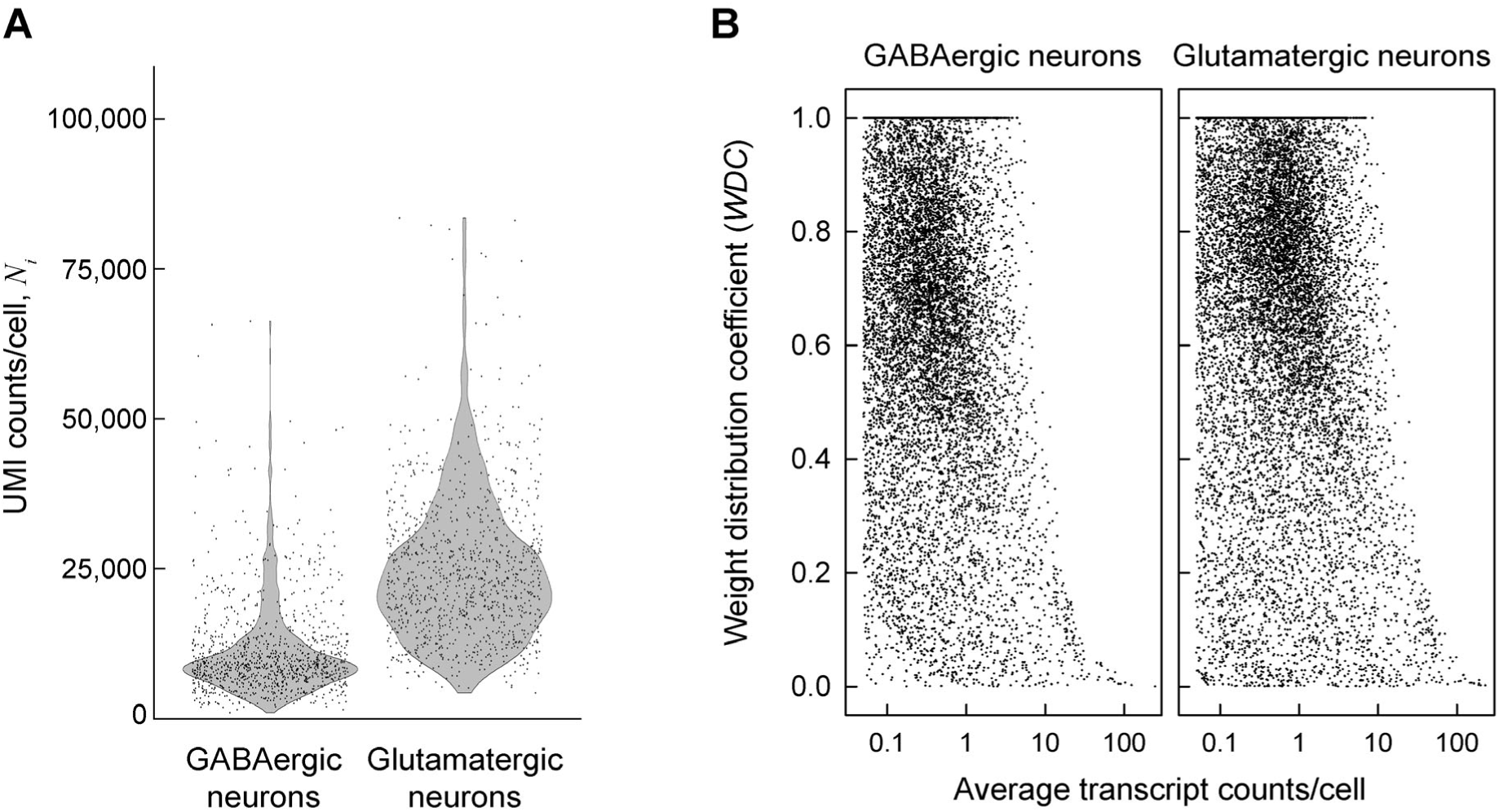
Measured statistical weights of cells in E18 mouse brain GABAergic and glutamatergic neurons do not match assumptions of common DGE analysis models. **A**. Total transcript (UMI) counts per cell *N_i_* in a combined E18 mouse brain dataset from 10X Genomics (see Methods). Each data point represents an individual cell. B. Weight distribution coefficient (*WDC*) for the statistical weight of different mRNA transcripts. Each point shows *WDC*_Z_ for a gene *Z* calculated directly from the measured transcript counts as described in Supplement 1. Standard, unweighted averaging of transcript counts utilized in most DGE analysis models assumes *WDC*_Z_ = 0. Utilization of aggregate counts in pseudo-bulk models assumes *WDC*_Z_ = 1. Neither reflects the reality.

The large variation in *N_i_*, however, necessitates not only normalizing the counts but also accounting for the statistical weight *w_Z,i_* of each cell *i* when calculating average relative expression 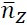. In general,

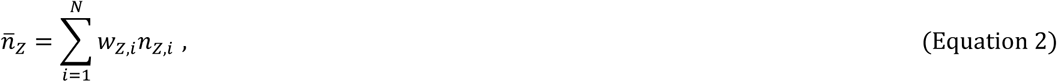

where ∑_i_*w_Z,i_* = 1 and *N* is the number of cells, as has been well established in statistics of experiments with non-uniform accuracy of data measurement ^32^.

In scRNASeq, *w_Z,i_* accounts for measurement precision of gene *Z* counts in cell *i* and therefore may be gene dependent. The most accurate unbiased estimate of the mean value (here 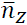) for a random variable (*n_Z,i_*) is obtained when *w_Z,i_* is proportional to the inverse variance of the variable ^32^, 1⁄*Var*(*n_Z,i_*). Highly variable *N_i_* leads to *w_Z,i_* that change from cell to cell and gene to gene.

The value of *w_Z,i_* can be calculated directly from the experimental data without any additional assumptions as

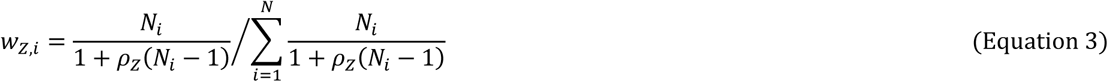

where 0 ≤ ρ_Z_ ≤ 1 is the intracluster correlation coefficient (ICC) calculated from the data (Supplement 1).

Equation 3 and several methods of calculating ρ_Z_ are well-known in the theory of cluster-randomized experiments, e.g., cluster-randomized clinical trials ^28,29^. To illustrate the concept, consider reporting of deaths in a drug trial. Death rates *n_Z,i_* may vary significantly between participant clusters at different trial sites *i*. These rates are more reliable in clusters with thousands of participants (e.g., large cities) compared to dozens of participants (e.g., small towns). They may be affected not only by the cluster size (*N_i_*) but also by other factors, e.g., differences in the lifestyle and healthcare between the clusters.

Correlations in the latter factors within each cluster necessitate weighing the counts based on Equations 2,3 ^28,29^. In the absence of differences in these factors, the same death rate is expected in all clusters and ρ_Z_ = 0. When ρ_Z_ = 0, *w_Z,i_* is proportional to *N_i_* (*w_Z,i_* = *N_i_*⁄∑_i_ *N_i_*), thus aggregating counts *N_Z,i_* from all clusters and yielding 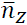 = ∑_i_ *N_Z,i_*⁄∑_i_ *N_i_* in Equation 2. At very strong correlations (ρ_Z_ = 1), the averaging of cluster death rates *n_Z,i_* becomes unweighted (equally weighted with all *w_Z,i_* = 1⁄*N*).

Generally, ρ_Z_ balances effects of the true outcome variability across clusters and imprecisions in measuring/estimating the outcome (relative counts such as death rate) in each cluster. When ρ_Z_ ≠ 0, measurement imprecisions become negligible in large clusters with ρ_Z_*N_i_* ≫ 1, all of which then have approximately the same weight that is larger than the weights of smaller clusters. At larger ρ_Z_, more clusters are sufficiently large to be weighed equally.

Supplement 1 shows that *n_Z,i_* averaging in scRNASeq is conceptually similar and describes how ρ_Z_ can be calculated from scRNASeq data without model assumptions. In scRNASeq, ρ_Z_ balances effects of the true biological variance between the cells and imprecisions in measuring *n_Z,i_*. The importance of weighing clinical trial results and accounting for the ICC has been recognized decades ago. We argue that the scRNASeq data analysis should recognize the importance of this approach as well.

Instead, almost all common approaches to scRNASeq data analysis use the unweighted averaging (AV), *w_Z,i_* = *w*_AV_ = 1⁄*N*, implicitly neglecting variable precision of the transcript count measurement from cell to cell (ρ_Z_ = 1 assumption). Pseudo-bulk approaches to data analysis from multiple samples aggregate the counts (AC), *w_Z,i_* = *w*_AC_ = *N_i_*⁄∑_i_ *N_i_*, implicitly neglecting correlated transcript expression within each cell and thereby expected biological variance in transcript counts between different cells (ρ_Z_ = 0 assumption). Statistical implications of such assumptions have been extensively studied in the context of cluster-randomized experiments. Both are known to cause false findings when the cluster size and composition are highly variable ^28–31^, which is the case in scRNASeq.

To examine implications of these assumptions in a real scRNASeq experiment, we selected cells expressing *Gad1*, *Gad2*, and *Slc6a1* (markers of GABAergic neurons) and cells expressing *Gls*, *Grin2b*, and *Slc17a7* (markers of glutamatergic neurons) from E18 mouse brain scRNASeq data from 10X Genomics (see Methods). These genes are not the only markers for the corresponding neurons, yet they are highly expressed and separate two minimally overlapping (< 1%), well-defined, and sufficiently large (∼ 1,000 cells each) subsets of data (Fig. 1A). Without necessarily requiring biological precision or accuracy, this was all we needed to test different DGE analysis models.

Instead of assuming *w_Z,i_* = *w*_AV_ (ρ_Z_ = 1) or *w_Z,i_* = *w*_AC_ (ρ_Z_ = 0), we evaluated ρ_Z_ and the corresponding *w_Z,i_* from measured *n_Z,i_* (Supplement 1) and calculated the weight distribution coefficient *WDC* defined as normalized variance of *w_Z,i_* (*WDC*_Z_ = *Var*(*w_Z,i_*)⁄ *Var*(*w*_AC_)). At *w_Z,i_* = *w*_AV_, *WDC*_Z_=0. At *w_Z,i_* = *w*_AC_, *WDC*_Z_=1. *WDC* values calculated from the data provide useful information about cell-to-cell variability in gene expression and reveal that neither of the two approximations is accurate for most genes (Fig. 1B).

*WDC*_Z_ ≈ 0 means that *w_Z,i_* is largely unaffected by the sequencing depth, indicating that the expression of the gene is variable enough for the transcript sampling error to be small compared to the gene expression variability. This is observed for the most highly expressed genes due to the low sampling error (Fig. 1B) and may also be expected for highly variable genes with moderate expression (e.g., genes regulating cell response to stress and environment). *WDC*_Z_ ≈ 1 means dominant contribution of the sequencing depth to *w_Z,i_*, indicating low expression variability from cell to cell and/or low expression of the gene. This may be expected, e.g., for tightly regulated cell maintenance genes with low-to-moderate expression. At the same expression level, lower *WDC* indicates higher expression variability.

The values of *w_Z,i_* significantly affect the fold-change (FC) in the average gene expression, which is the reported measure of DGE. For instance, log_2_(FC) between glutamatergic and GABAergic neurons is significantly different for many genes when calculated with measured *w_Z,i_* rather than assuming *w_Z,i_* = *w*_AV_ (Fig. 2A). For some genes, this changes even the sign of log_2_(FC) so that upregulated genes may appear to be downregulated and vice versa. The difference between weighted and unweighted FC can be more than 2-fold and have dramatic implications for data interpretation when the genes of interest are affected, even though weighted and unweighted log_2_(FC) values are close for many genes.

**Figure 2.**
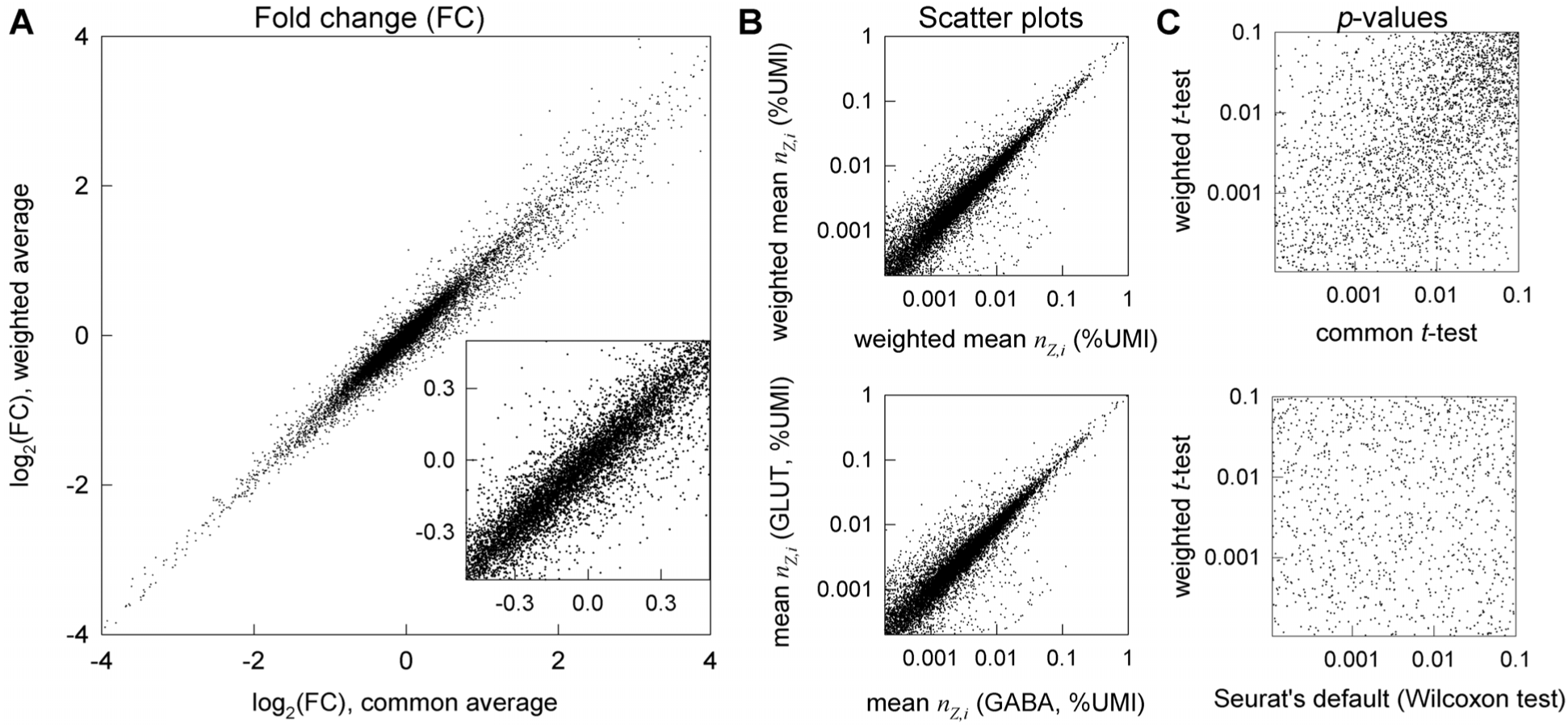
Statistical weights of cells significantly affect the calculated log_2_(FC) and *p*-values. **A.** Effects of the statistical weights on log_2_(FC) between glutamatergic vs. GABAergic neurons in E18 mouse brain (same as in Fig. 1A) plotted for 12,087 genes with at least 50 total counts in glutamatergic neurons. 7% of these genes (847) have different signs of log_2_(FC) for common and weighted averages. 17% of the latter (143) have the difference between the two log_2_(FC) values > 0.25 and 3% (29) larger than 0.5. **B.** Scatter plots for mean (bottom) and weighted mean (top) expression of these genes in GABAergic (x axis) and glutamatergic (y axis) neurons. **C.** Effects of the statistical weights on *p*-values for genes plotted in **A**. The common (unweighted) *t*-test was performed based on relative count rather than logarithmic normalization to avoid artifacts of the latter. Among the genes with at least 50 total counts in glutamatergic neurons, the weighted *t*-test found 3,993 genes with *p*<10^-4^ (of which 6 were not found by the common *t*-test with *p*<0.05 and 845 were not found by the Seurat’s default Wilcoxon test with *p*<0.05. The common *t*-test found 3,747 genes with *p*<10^-4^ (of which 3 were not found by the weighted *t*-test with *p*<0.05). The Wilcoxon test found 8,045 genes with *p*<10^-4^ (of which 3,523 were not found by the weighted *t*-test with *p*<0.05), which was clearly inconsistent with the scatter plots for either mean or weighted mean expression.

### Weighted statistical testing describes DGE significance without model assumptions

Statistical weight values affect not only the log_2_(FC) value but also statistical significance of DGE (the probability *p* of the observed log_2_(FC) when the expected log_2_(FC) is zero). To account for this, we cannot use common approaches to calculate the *p*-values in scRNASeq. Not only do these approaches use assumed weights but they also rely on additional assumptions, sometimes implicit.

For instance, the *p*-values are commonly obtained from the Wilcoxon test (default in Seurat), in which all data points are ranked (1,2,3…) in the order of increasing *n_Z,i_*. This test is appropriate and useful when the question is whether the distributions rather than mean values of random variables are different ^33^. However, its *p*-value is not the probability of the observed log_2_(FC) when the expected one is zero ^33^. The Wilcoxon test may return low *p* values more often than expected when cell-to-cell distributions of transcript counts in two samples are different even when the relative expression is the same (log_2_(FC) = 0). For instance, scatter plots indicated similar relative expression of most genes in glutamatergic and GABAergic neurons (Fig. 2B). In contrast, the Wilcoxon test found over 60% of genes being differentially expressed with *p* < 10^-4^ (Fig. 2C), in part because of very different total number of transcripts per cell in these neurons (Fig. 1A). The variation in total (and consequently individual gene) transcript distribution is common in scRNASeq because of different transcriptional activity of the cells, different sequencing depth (detected fraction of transcripts), and effects of sample preparation. Similar details must be considered when applying common *t*-test on log-normalized data in Seurat. The test compares the mean values of log-normalized counts (logarithms of scaled relative counts plus a pseudo count.) This differs from comparing the mean values of relative counts, and we show the consequences in the next Section.

Rather than assuming any properties of transcript distribution explicitly or implicitly, we calculate the *p*-values directly from raw data relying only on sufficient numbers of relevant cells (from ∼ 100 to thousands of cells per group is becoming common and likely to increase). Mean values in such large datasets are expected to have normal distributions (central limit theorem ^33^). Therefore, the corresponding *p*-values can be found from a *t*-test, a weighted version of which accounts for variable statistical weights (Supplement 1). This approach is similar to statistical testing of weighted averages in cluster-randomized experiments, which has been used for many years ^28,29,31^. For instance, the *p*-values for DGE between glutamatergic and GABAergic neurons calculated from the weighted *t*-test are very different from those calculated from the Wilcoxon test or the common (unweighted) *t*-test (Fig. 2C). Almost half of the genes with highly significant DGE (*p* < 10^-4^) found by the Wilcoxon test are not differentially expressed in the weighted or unweighted *t*-test (*p* ≥ 0.05, Fig. 2C), consistent with the scatter plots for the relative gene expression (Fig. 2B).

It is important to keep in mind, however, that the weighted *t*-test treats individual cells as independent experimental replicates (like all common approaches to scRNASeq analysis). Depending on the experiment and cell type identification algorithm, this may not be a good approximation (Supplement 1). When multiple sets of experimental data are available, each independent set can be used as a biological replicate to resolve this problem. When such replicates are not available, answering a different statistical question with a supplemental χ^2^ test may be useful for reducing the resulting false positive findings.

The χ^2^ test compares aggregate transcript counts just in the sequenced cells rather than average counts per cell in the specimens from which the cells have been sampled (Supplement 1). Because it accounts only for variance in stochastic transcript sampling but not for variance in the sampling of cells or variance in gene expression from cell to cell, it may significantly underestimate the total variance and should not be used alone. However, it may mitigate cell sampling bias effects by excluding the weighted *t*-test findings based on χ^2^ test *p*-values larger than a minimum significance threshold, e.g., 0.05.

To illustrate this and other approaches to combining the weighted *t*-test with the χ^2^ test, consider DGE between GABAergic and glutamatergic neurons selected for Fig. 2. For these cells, the weighted *t*-test detected 3,993 genes with *p* < 10^-4^, among which 3 genes had χ^2^-test *p* ≥ 0.05 (no significant difference in the aggregate counts). The highly significant weighted *t*-test for the latter genes was therefore caused by a difference in the transcript distribution among the cells rather than a difference in the aggregate gene expression. This may be a real biological difference between the two types of neurons or just an effect of biased cell sampling from mouse brain. The best solution for this dilemma is replicate or independent experiments, eliminating the need for the χ^2^-test. A moderately conservative proxy approach when such experimental data are not available is to use the *p*-value from the χ^2^-test instead of the weighted *t*-test one when the former but not the latter exceeds the minimum significance threshold. This is the approach utilized hereafter. One may also just select the larger of the two *p-*values to minimize false positive findings, which is the most conservative approach to combining the tests.

We found the DGE results based on the weighted-*t*-test/χ^2^-test combination to be very different from those produced by standard DGE analysis approaches. In the same glutamatergic vs. GABAergic neurons example from Fig. 2, the Wilcoxon test detected 3,629 genes with *p* < 10^-4^ that were not significant (*p* > 0.05) in the weighted-*t*-test/χ^2^-test combination (note that Fig. 2 presents analysis based on the weighted *t*-test alone). The latter approach detected 845 genes with *p* < 10^-4^ that were not significant in the Wilcoxon test. Among these 845 genes, 147 had more than 2-fold difference in expression. For instance, the Wilcoxon test did not detect a change in expression of cancer-associated genes *Tacc3* and *Birc5*, which had more than 0.2 counts/cell in both groups of neurons, more than 3-fold change in expression, and FDR-adjusted *p* < 10^-5^ in the weighted-*t*-test/χ^2^-test combination. Other tests built into Seurat’s FindMarkers function had similarly large discrepancies with the weighted-*t*-test/χ^2^-test combination when the logarithmic data normalization recommended by Seurat was used. The discrepancy with the unweighted *t*-test was less dramatic when relative counts normalization was used, yet it was still significant (Fig. 2B). For instance, the unweighted *t*-test found 21 genes with *p* < 10^-4^ that had *p* ≥ 0.05 in the weighted-*t*-test/χ^2^-test combination.

### Weighted statistical testing improves robustness and sensitivity of DGE analysis

This analysis suggests that traditional approaches may produce misleading results, yet it does not prove their failure. Comparison of experimental data reveals major differences between model predictions but cannot establish which model is correct since the ground truth in these experiments is unknown. It is therefore also important to test predictions of different models by using simulated data that are based on real-world experiments yet model well defined data patterns, in which the ground truth is known.

First, we tested the simplest pattern of UMI subsampling in glutamatergic neurons (Fig. 3A-G). To generate this pattern, we used glutamatergic neuron data, in which we randomly subsampled all UMIs for all genes with 40% probability of retaining each UMI. Such 40% subsampling mimics reduced sequencing depth (60% fewer counts detected by sequencing), which is well within normal experimental variation. We performed 10 random subsampling simulations, using only genes with more than 30 total counts in the original data and analyzed DGE effects of the subsampling. Since random subsampling does not change relative gene expression, any DGE finding in this simulation is a false positive one. This simulation revealed dramatic failure of all commonly used DGE analysis models (Fig. 3C,D,F,G). The false positive DGE was still unacceptable at 60% subsampling (Supp. Fig. S2.1).

**Figure 3.**
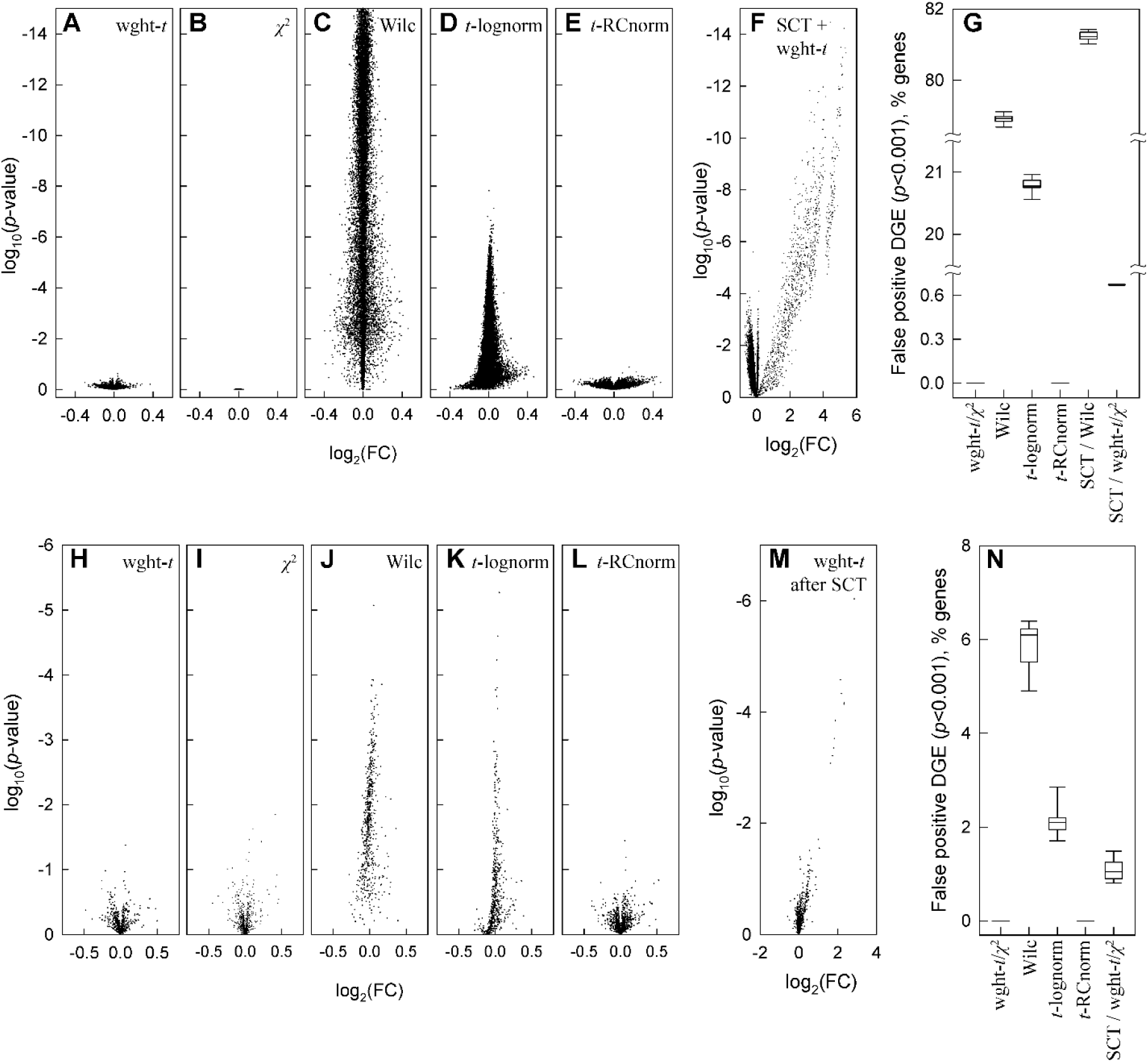
Weighted statistical testing eliminates false positive DGE in analysis of original vs. subsampled glutamatergic neurons data. **A-G**. False DGE discovery after SCTransform (SCT) or after 40% count subsampling (retaining 40% of counts for each gene by randomly selecting counts to be removed). 12,911 genes are shown, each with more than 30 counts in all cells in the original data. **A-E.** Volcano plots for a single subsampling simulation after weighted *t*-test (**A**), χ^2^-test (**B**), Seurat’s FindMarkers function default Wilcoxon test (**C**), Seurat’s FindMarkers optional t-test for logarithmically normalized data (**D**), and *t*-test for RC normalized data (**E**, Equation 1). **F**. DGE artificially introduced into the data by count rescaling with SCT, as revealed by weighted *t*-test comparison of the data before and after SCT. **G.** Fraction of the 12,911 genes (%) falsely discovered as differentially expressed with *p* < 0.001 after 10 subsampling simulations. **H-N.** DGE analysis for 1,000 randomly selected genes (each with more than 30 total counts in all cells) altered by random, uniformly distributed noise. **H-L.** Volcano plots for the same tests as in **A-E**. **M.** Volcano plot for weighted *t*-test comparison preceded by SCT of both original and noise-altered data. **N.** Fraction the 1,000 noise-altered genes falsely discovered as differentially expressed with *p* < 0.001 after 10 simulations. In **G** and **N**, boxes show 25th to 75th percentile range, whiskers show 10th to 90th percentile range, and lines show median values.

The weighted *t*-test, χ^2^ test, and their combination had no false positive findings in any of the 10 simulations even at *p* < 0.05 cutoff (Fig. 3A,B). In contrast, Seurat’s FindMarkers function with default Wilcoxon test consistently identified over 75% of all genes as differentially expressed with *p* < 0.001 and over 60% as differentially expressed with *p* < 10^-5^ (Fig. 3C,G). Seurat’s FindMarkers function with the common *t*-test option consistently identified over 20% of all genes with *p* < 0.001 and > 1% with *p* < 10^-5^.

Over 1,000 genes with highly significant false DGE discovered by the default Seurat’s pipeline for DGE analysis had |log_2_(FC)| > 0.1 (the default threshold in Seurat) and dozens had |log_2_(FC)| > 0.3 (as reported by Seurat). The results for other tests built into Seurat were qualitatively similar. When the commonly recommended logarithmic normalization (default in Seurat) was replaced with non-parametric RC (relative count) one (Equation 1), the unweighted *t*-test did not identify DGE (Fig. 3E,G), revealing unacceptable artifacts of logarithmic normalization (which affects the *t*-test but not the Wilcoxon test).

The main cause of the dramatic failure of standard DGE analysis models was count dropout (loss of some nonzero counts due to subsampling). This is a well-known effect of sample-to-sample variations in sequencing depth and/or RNA recovery from cells, which is one of the reasons behind model-dependent data regression and/or rescaling ^34–37^. RC normalization combined with the weighted-*t*-test/χ^2^-test combination or the unweighted *t*-test make such model-dependent data processing unnecessary.

We found that SCTransform (SCT) – one of widely used data rescaling procedures – was inefficient in reducing DGE caused by UMI subsampling and produced its own artifacts. SCT, which rescales counts based on negative binomial regression, was developed in part to reduce the count dropout effects ^7^. We tested how SCT version 2 with default parameters affected DGE analysis in Seurat. Even the most conservative weighted-*t*-test revealed false DGE between original data before and after SCT, i.e. SCT introduced artificial expression patterns absent from the data (Fig. 3F). SCT increased rather than reduced artifacts of count dropouts, e.g., increasing false positive DGE discovered by the Wilcoxon test (Fig. 3G). Moreover, we observed false DGE when SCT was used for original and subsampled data followed by the weighted-*t*-test/χ^2^-test combination (Fig. 3G). SCT artifacts were particularly strong for low expression genes with 30-50 counts per 1,000 cells and gradually disappeared at higher expression.

Next, we altered a subset of 1,000 randomly selected genes in the glutamatergic neuron gene count matrix *N_Z,i_* (the dataset in Fig. 2A) by random, uniformly distributed noise. Again, only genes with more than 30 total counts in all cells were included. The added noise had zero average amplitude, maximum amplitude between 0.8*N_Z,i_* and *N_Z,i_* at *N_Z,i_* ≤ 5, and variance 5*N_Z,i_* at large *N_Z,i_* (see Methods). 10 random sets of 1,000 genes were tested. The noise was randomly regenerated for each simulation.

Again, the weighted-*t*-test/χ^2^-test combination and common *t*-test after RC normalization found no false positive DGE (Fig 3H,I,L,N). The other tests discovered DGE for many more genes than expected by chance at the corresponding *p*-values (Fig. 3J,K,M,N). Like in the case of count subsampling, SCTransform produced rather than removed false DGE. Because the simulated noise produced fewer count dropouts than the count subsampling, its effects were less dramatic yet still significant. Overall, the effects of random noise and count subsampling were very similar.

We then tested how well different models discover DGE masked by noise and tested whether SCT helped the best performing model to suppress noise and reveal DGE. For these simulations, we combined the same random noise as above with a proportional increase (50%) in the counts of a 300 gene subset of the 1,000 genes modified by noise.

The weighted-*t*-test/χ^2^-test combination was by far the most accurate and reliable (Fig. 4). Its benefits were most apparent for moderate expression genes (average 0.3 to 3 counts per cell), in which the added noise was comparable or exceeding the added DGE signal. It missed significantly fewer upregulated genes compared to other tests (Fig. 4A,C) and did not detect any false DGE (Fig. 4B,D). The *t*-test for RC normalized data did not detect false DGE either, yet it missed more upregulated genes. The Wilcoxon test and *t*-test for log-normalized data reported significant false DGE.

**Figure 4.**
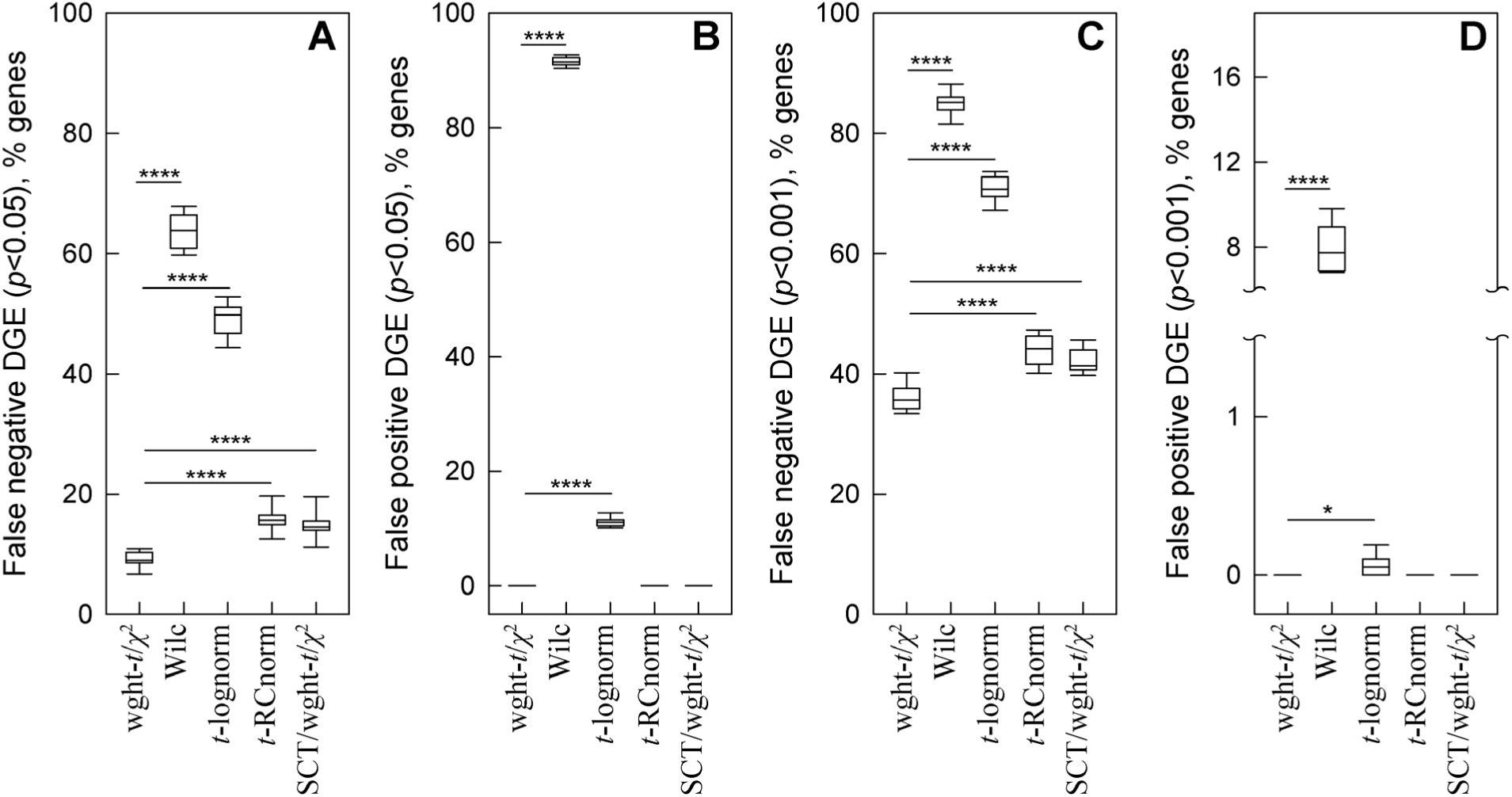
Weighted statistical testing reduces false negative and false positive DGE in simulations of gene upregulation superimposed with comparable noise. Random subsets of moderately expressed genes (0.3 to 3 counts per cell) in glutamatergic neurons were selected for the simulations. **A.** False negative DGE (*p* >0.05 and/or log_2_(FC) < 0.1) among 300 genes modified by 50% increase in expression and comparable random noise. Weighted-*t*-test/χ^2^-test combination was used without SCT (wght-*t*/χ^2^) and with SCT (SCT/ wght-*t*/χ^2^). Wilcoxon test (Wilc), *t*-test for log-normalized data (*t*-lognorm), and *t*-test for RC normalized data (*t*-RCnorm) were used without SCT. **B.** False positive DGE (*p* < 0.05 and log_2_(FC) > 0.1) among 1,000 genes modified only by random noise. **C** and **D**. Same as **A** and **B**, respectively but at *p* < 0.001 threshold. The box and whiskers parameters were the same as in Fig. 3G; **** means *p* < 0.0001 and * means *p* < 0.05.

In moderately expressed genes, SCT did not cause detection of false DGE, but it reduced the fraction of upregulated genes detected by the weighted-*t*-test/χ^2^-test combination (Fig. 4A). In genes with average < 0.3 counts/cell, the DGE signal was fully masked by noise and not detected by any tests. In genes with average > 3 counts/cell, the DGE signal was stronger than the noise and reliably detected by all tests.

Finally, we confirmed the weighted *t*-test performance by analysis of simulated data with predefined data distributions (Supplement 1, Fig. S1.4). Since all in silico cells were independent replicates, in these simulations we used only the weighted *t*-test but not its combination with the χ^2^-test. The weighted *t*-test had the best combination of power for discovering true DGE (low false negative findings) with type I error rate control (low false positive findings). The Wilcoxon test had poor type I error rate control when total numbers of transcripts per cell in the two samples were different (an analogue of different sequencing depth). The unweighted *t*-test was too conservative and had significantly higher false negative findings.

### Weighted statistical testing outperforms the pseudo-bulk approach

When scRNASeq data is available from multiple samples per genotype or treatment, the biological variability and differences in measurement accuracy between the samples become important. The pseudo-bulk (PB) approach, which has been developed and is commonly used for analysis of such experiments ^1,38^, however, involves several questionable and unnecessary simplifying approximations.

First, the PB approach uses count aggregation within each sample instead of weighted averaging, implicitly assuming *w_Z,i_* = *w*_AC_ = *N_i_*⁄∑_i_ *N_i_*. As discussed above, this approximation may work well for some genes, yet it may significantly affect the results for many other genes (Fig. 1B).

Second, the PB approach neglects sample-to-sample variations in measurement accuracy. This accuracy can be characterized by the variance of 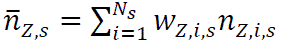 for each sample *s*. To be internally consistent, the aggregated count analysis must use weighted average of 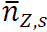 with statistical weights dependent on this variance ^32^, which can be calculated from the experimental data as described in Supplement 1.

Third, the PB approach determines the variances for statistical testing from negative binomial regression (e.g., using DESeq2 ^39^ for identifying DGE in Seurat). Instead, these variances can be calculated directly from the data without any approximations as described in Supplement 1.

To avoid these approximations, one can use *n_Z,i_* measured in each scRNASeq experiment to determine the experimental values of *w_Z,i_* as described above followed by calculating the corresponding 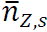 and *Var*(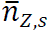). The log_2_(FC) and *p* values can then be calculated by using the weighted *t*-test with weights determined from the experimental data as described in Supplement 1. Although 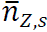 distribution may not be normal, the *t*-test is still the most robust approach when only small number of replicates are available, and the statistical distribution cannot be reliably established ^40^. This is almost always the case in high cost scRNASeq experiments.

Note that a PB approach incorporating weights was also described, but these weights were based on fitting logarithmically normalized data to a generalized linear model rather than determined directly from the data without model assumptions.^41,42^

To test our minimal assumptions approach, we again simulated datasets with the same well-defined noise and DGE signal based on the same glutamatergic neuron data. We generated 10 samples by randomly subsampling the cells from the original dataset. To mimic effects of variable sample size, the samples contained 20%, 40%, 60%, 80%, or 100% of the neurons. In 5 of the samples, 1,000 moderate expression genes (average 0.3 to 3 counts per cell) were modified by random noise. In the other 5 samples, the same 1,000 genes were modified by similar random noise and by an increase in expression of a 300 gene subset. The noise-only and noise+DGE signal modified samples were then compared using the weighted *t*-test for weighted average counts (Fig. 5, wght-*t)*, unweighted *t*-test for aggregate counts (Fig. 5, Aggr/*t*-test), weighted *t*-test for aggregate counts (Fig. 5, Aggr/wght-*t*), and DESeq2 test for aggregate counts (Fig. 5, PB/DESeq2, Seurat’s default for traditional PB analysis).

**Figure 5.**
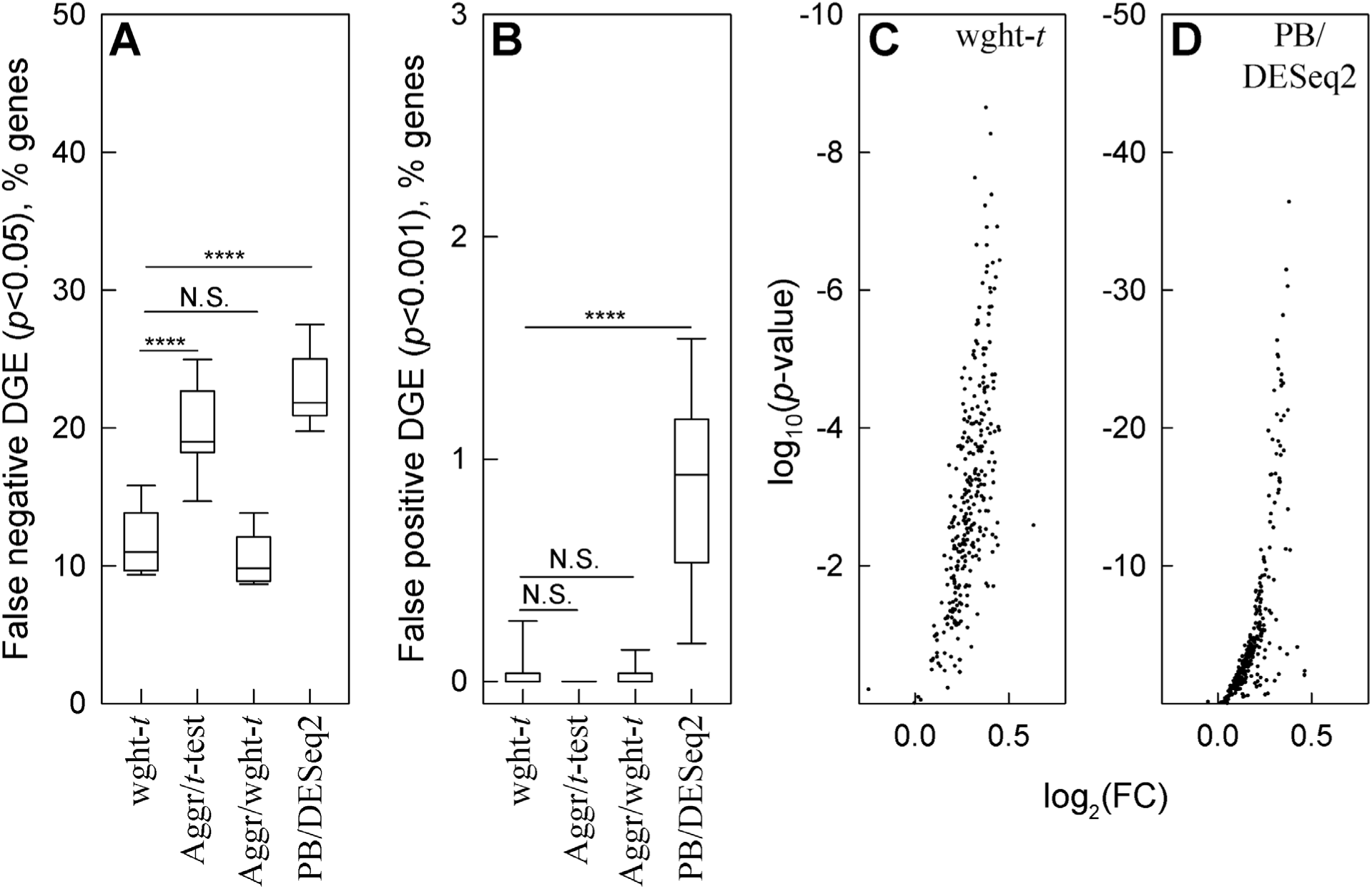
Weighted statistical testing outperforms the pseudo-bulk approach to DGE analysis. Two 5-sample sets were generated from glutamatergic neuron data by random cell subsetting. Like in Fig. 4, we added random noise to a 1,000 gene set that had 0.3 to 3 counts/cell and 30% increase in expression to a 300 gene subset. **A.** False negative DGE among 300 genes with increased expression (wght-*t* is the weighted *t*-test for weighted average counts, Aggr/*t*-test and Aggr/wght-*t* are unweighted and weighted *t*-tests for aggregate counts, PB/DESeq2 is pseudo-bulk aggregate count analysis with DESeq2). **B.** False positive DGE among 700 genes modified only by noise. **C** and **D**. Volcano plots for the genes modified by the expression increase and noise analyzed by weighted *t*-test (**C**) and PB/DESeq2 (**D**). In panels **A** and **B**, **** means *p* < 0.0001, *** – *p* < 0.001, * – *p* < 0.05, N.S. – not significant.

Since we found multiple sample analysis of aggregate counts to be less sensitive to noise and more sensitive to changes in expression, we made the comparison more stringent than in Fig. 4 and increased the expression by only 30%. We also selected only 500 from the 1,000 modified genes randomly. We selected the other 500 as the genes with the largest cell-to-cell expression variability (coefficient of variation), while we still selected a 300 gene subset of the 1,000 genes for expression upregulation randomly.

These simulations supported our theoretical deductions. Weighted *t*-test again detected the largest fraction of upregulated genes, and weighted averaging resulted in fewer false negative findings than count aggregation (Fig. 5A). Weighted averaging and count aggregation produced similar results in this simulation, probably because count aggregation worked well for lower expression genes used in it (Fig. 1B). Both weighted and unweighted *t*-tests for relative counts still had low false positive findings (Fig. 5B). DESeq2 analysis of aggregate counts ^39,43^ – the Seurat’s default and widely used PB method – had the largest number of false negative (Fig. 5A) and false positive (Fig. 5B) findings. As expected ^1^, this PB analysis performed better than the Wilcoxon test (c.f., Fig. 4), yet it still had too many false findings (e.g., 1% of ∼20,000 genes detected by modern assays would result in ∼200 false positive findings). Moreover, it’s up to 50 orders of magnitude lower *p*-values compared to the weighted *t*-test indicated severe variance underestimation (Fig. 5C,D). When we simulated DGE ideally matching DESeq2 assumptions (equal number of 150 up– and 150 down-regulated genes), the false positive findings disappeared yet the false negative findings remained high (Supp. Fig. S2.2).

### Weighted statistical testing reduces false positive findings in spatial transcriptomics

The approach to DGE analysis described above can also be used for spRNASeq applications after several small modifications. The spRNASeq data is a matrix *N_Z,i_* of gene Z counts in a spatial bin *i*. In some assays (e.g., Visium HD™ from 10X Genomics), the spatial bins are fixed areas like squares. In other assays (e.g., Xenium™ from 10X Genomics), the bins are segmented areas designed to represent individual cells. From the perspective of DGE analysis, the main differences between scRNASeq and spRNASeq are: (a) significantly fewer transcript counts per bin, (b) overlaps of different cells within the same bin (even after cell-based segmentation), and (c) assay noise correlations between adjacent bins or cells due to RNA probe diffusion.

Because of these differences, the aggregated count approximation with *w_Z,i_* = *w*_AC_ = *N_i_*⁄∑_i_ *N_i_* (ρ_Z_ = 0) may be the best one for the currently available assays (Supplement 1). In full transcriptome assays like Visium HD, log_2_(FC) can then be calculated from the corresponding weighted averages and the *p* values can be determined from the weighted-*t*-test/χ^2^-test combination. These log_2_(FC) and *p*-values must then be interpreted as the difference in aggregated rather than individual cell gene expression, which is the downside of the aggregated count approximation. An alternative solution – enabling one to account for biological variability between cells of the same type within each sample – is to define the sample as a group of multiple cell clusters with each cluster containing multiple bins (see Supplement 1).

For instance, in a Visium HD assay, mRNA within a tissue section is hybridized with gene-barcoded probes, which are then transferred onto a slide coated with UMI and spatially barcoded primers. The spatial barcode distinguishes 2×2 µm squares on the slide surface. After probe hybridization with the primers, a cDNA library is constructed, amplified by PCR, sequenced by next generation sequencing like in scRNASeq, and converted into an *N_Z,i_* matrix of gene *Z* count in spatial bin *i*. A user may select the number of 2×2 µm squares included in each spatial bin. Our testing suggests 10-15 µm smearing of the probes due to diffusion when they are transferred from tissue section onto a Visium HD slide. We therefore agree with the 10X Genomics recommendation of using 8×8 µm binning.

The *N_Z,i_* data matrix generated by Visium HD can be analyzed in the same way as in scRNASeq. The values of log_2_(FC) and *p* comparing two groups of bins can be calculated by setting ρ_Z_ = 0 and combining the weighted-*t*-test and χ^2^-test. In this case, the *p*-values predicted by the weighted *t*-test and χ^2^-test may be similar. At lower counts per bin/cell, the variance associated with random transcript sampling for NGS may be dominant, in which case the weighted *t*-test and χ^2^-test p-values are expected to be close to each other (Supplement 1). We recommend using the more conservative *p*-value estimate from the two tests. Groups of bins preferentially or exclusively containing cell type(s) of interest may be selected for such comparison as illustrated by Fig. 6A for proliferating chondrocytes in the growth plate. Alternatively, bins may be grouped or clustered based on gene expression patterns. Analysis of data with multiple samples per genotype or treatment again follows the protocol described above.

**Figure 6.**
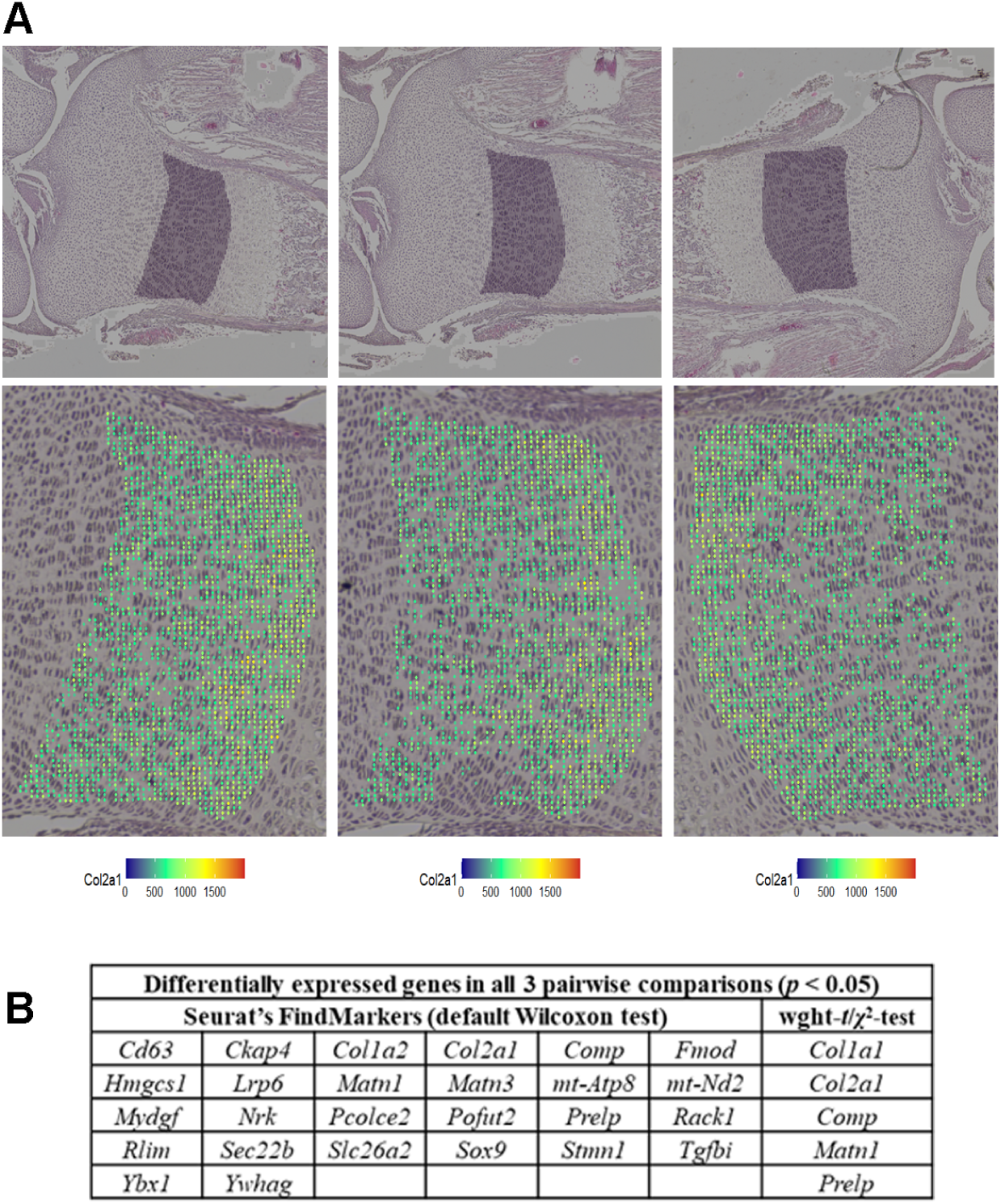
Weighted statistical testing reduces false positive DGE in sp-RNASeq (Visium HD assay). Top panels in. **A.** Proliferating chondrocytes were selected in 3 sections from the same E18.5 tibia. Shaded areas show the selected regions in high resolution images of H&E-stained sections. **Bottom panels in A.** Zoomed images of the selected areas. Colored dots mark 8×8 µm bins used for the analysis of gene expression, which passed quality control based on the following criteria: > 20 features per bin, > 100 counts other than *Col1a1* or *Col1a2* per bin, mitochondrial genes between 0.1 and 5% UMI, and *Col2a1* > 5% UMI. The bin color represents the relative expression of *Col2a1* (*n*_Col2a1,i_ × 10,000). **B.** DGE analysis performed with the weighted-*t*-test/χ^2^-test combination and Seurat’s FindMarkers function for all 3 possible pairwise comparisons between the selected areas. Only genes detected in at least 10% of the bins were analyzed.

Note that accounting for statistical weight is even more important for spatial than single cell RNASeq. Large (∼ 100-fold) variations in the statical weight of bins cannot be eliminated even in a hypothetical perfect spatial assay because of large lateral variations in RNA density in tissue sections, which are biological rather than technical. Figure 6B illustrates their effects based on comparing proliferating (and possibly some pre-hypertrophic) chondrocytes in 3 sections from the same growth plate. Our approach identified differential expression only in several extracellular matrix genes, expression of which may vary locally within the same growth plate. In contrast, the analysis based on Seurat’s FindMarkers function identified many additional genes not expected to be differentially expressed within the same growth plate. It was the attempt to understand this failure of Seurat’s traditional analysis that initiated this whole study.

Importantly, analysis of imaging rather than sequencing based spatial transcriptomics assays (e.g., Xenium) requires a different approach to data normalization. These assays image signals from probes for a limited preselected subset of genes. They also cannot identify more that several total transcripts per µm^2^ because of limited optical resolution. Moreover, ratios of counts between different gene transcripts appear to change from experiment to experiment (at least in our experience of using the Xenium assay). Together, these assay features preclude relative count normalization and utilization of statistical approaches to data analysis described above. We have found several alternative solutions, but their discussion is beyond the scope of this paper.

## DISCUSSION

The key features distinguishing our approach to sc– and sp-RNASeq analysis from commonly used ones are: (1). No assumptions about noise statistics besides randomness. (2) Model-free noise averaging rather than model-dependent noise regression. (3). Unbiased accounting for technical noise variations from cell to cell and sample to sample based on statistical weights deduced directly from the data.

Fewer assumptions inherently produce less biased outcome since data may not meet the assumptions. Using statistical weights reduces uncertainty in whether a gene is downregulated or upregulated (Fig. 2). Weighted statistical tests reduce the likelihood of interpreting random noise and variations in sequencing depth as DGE (Fig. 3). Together, accurate weighted averaging and statistical tests remove noise artifacts and detect DGE signals more reliably than data parametrization, rescaling, or noise regression (Figs. 3,4). They produce more consistent results when data from multiple samples are available as well (Fig. 5). Besides, the same approach with only minor adjustments produces more meaningful results for high-resolution spatial transcriptomics (Fig. 6).

To be beneficial, additional assumptions must accurately describe the assay noise. This does not seem to be the case for common scRNASeq analysis tools, as indicated by Fig. 1B and increased false findings by these tools (Figs. 3-6). One explanation is the noise associated with the cell-to-cell variation in the sequencing depth (counts per cell, Fig. 1A), which may be impossible to model (Supplement 1). Consider, e.g., the SCTransform (SCT) rescaling of transcript counts designed to correct the data for this variation ^7^. Fig. 3F shows a strong signal introduced by the SCT into the data. This signal is at least partially artificial rather than unmasked from the noise, which is indicated by the highly significant DGE between the original and randomly subsampled data after the SCT (Fig. 3G). The SCT artifacts are more pronounced for lower expression genes because of stronger sequencing depth effects on them. Together, Figs. 3F and 3G show that model-dependent tools like the SCT are not universally appropriate for scRNASeq. Therefore, we think that such tools should be avoided or at least tested for each data set. Random subsampling testing is essential because of the expected sequencing depth variation between samples.

While developing our minimal assumptions approach, we have made several observations that are worth keeping in mind when interpreting the results produced by our or any other DGE analysis:

1. Statistical weights of cells are not just mathematical constructions we used for data analysis; they are measures of gene count accuracy in each cell. Unweighted averaging may alter the apparent change in gene expression by more than a factor of 2 and affect even the sign of the change (Fig. 2A). While providing reasonable control of false positive findings, unweighted *t*-test for relative counts is too conservative and has less power in detecting DGE than the weighted *t*-test (Supplement 1, Fig. S1.4). Statistical weights are therefore important for data interpretation ^28,29,31,32^.
2. Violin plots of normalized transcript counts are visually appealing yet may be misleading even when one looks at the data points rather than the violins themselves. More graphically accurate representation of the same data by bona fide histograms may be equally misleading because histograms of normalized data do not account for statistical weights of cells. A peak due to cells with low confidence count values can be easily misinterpreted. For instance, violin plots or bona fide histograms may look very different for samples with identical gene expression when the sequencing depth is different (Fig. S1.5). Similarly, it is possible to imagine the situation when average gene expression is different in two samples, while the corresponding violin plots look the same. In contrast, histograms for unnormalized total transcript counts per cell faithfully represent the reality (Fig. 1A), because the statistical weights of cells are not important for their interpretation.
3. Analysis of DGE between cells selected based on cell clustering algorithms is questionable, unlike DGE analysis between cells selected based on known marker gene expression. This concern is recognized in Seurat documentation (FindMarkers function). However, being often ignored in practical applications, it may be an important contributing factor to false positive and false negative DGE results. All statistical models for *p*-value calculation presume random, independent sampling. All common cell clustering algorithms group cells based on differences in expression of thousands of highly variable genes (e.g., 2,000-3,000 genes in Seurat). The *p*-values calculated for these genes (and genes dependent on them) are questionable because cell clustering based on expression of these genes is inconsistent with independent sampling. Clustering based on several marker genes like in Fig. 1A and excluding these marker genes from DGE analysis resolves this concern. Strategies for identifying such marker genes when they are not known a priori are beyond the scope of the present work.
4. Analysis of false discovery rate (FDR) due to multiple comparison in scRNASeq is important ^44^ yet may be misused. Proper FDR analysis is question and context dependent. FDR is the fraction of genes with *p*-values below a predetermined threshold that is expected to be false positive by chance upon multiple comparisons ^44,45^. It is different for a specific group of genes, a pathway, or a list of candidate genes for follow up. However, FDR is often calculated for the full transcriptome, which is not always appropriate. Validating genes selected based on uncorrected *p*-values by additional or independent experiments reduces the risk of missing important genes by chance due to an FDR threshold, although it increases the study cost. With all this in mind, our scripts do not calculate corrected *p*-values, which can be easily determined, e.g., by using the p.adjust() function in R.

In conclusion, more accurate scRNASeq and spRNASeq data analysis with fewer assumptions will hopefully simplify identification of up/down-regulated genes and eventually reduce or eliminate the need for follow up. In the meantime, our findings suggest caution in interpreting unvalidated output of common DGE analysis approaches (Figs. 2-6). Since we used these approaches before, we reanalyzed our past experiments and learned useful lessons. Revisiting such data may be worth the invested time.

## RESOURCE AVAILABILITY

### Lead contact

Requests for further information and resources should be directed to and will be fulfilled by the lead contact, Sergey Leikin.

### Materials availability

This study did not generate new unique reagents.

### Data and code availability

- Spatial data have been deposited at NCBI GEO as https://www.ncbi.nlm.nih.gov/geo: GSE299816 and are publicly available.
- Additional data and simulations not included in this publication have been deposited at Synapse.org and are publicly available as of the date of publication at https://doi.org/10.7303/syn66520062.
- This paper also analyzes existing, publicly available data, accessible at 10X Genomics: https://www.10xgenomics.com/datasets/10-k-mouse-e-18-combined-cortex-hippocampus-and-subventricular-zone-cells-single-indexed-3-1-standard-4-0-0 and https://www.10xgenomics.com/datasets/10-k-mouse-e-18-combined-cortex-hippocampus-and-subventricular-zone-cells-dual-indexed-3-1-standard-4-0-0.
- All original code has been deposited at GitHub (https://github.com/sergeyleikin/sc-sp-RNASeq) and is publicly available.
- Any additional information required to reanalyze the data reported in this paper is available from the lead contact upon request.

## ACKNOWLEDGMENTS

This work was supported by the Intramural Research Program of NICHD. The Visium HD assay was performed with the assistance of Drs. James Iben and Fabio Rueda Faucz at the Molecular Genomics Core of NICHD. The authors thank Drs. Ryan Dale, Patrick Fletcher, Edward Mertz, and Shyamal Peddada for many useful discussions. A blog post by Jan Vanhove, https://janhove.github.io/posts/2016-02-16-cluster-randomisation-correction/ helped GM to realize the connection between our work and theory of cluster-randomized experiments.

## AUTHOR CONTRIBUTIONS

Conceptualization, G.M. and S.L.; methodology, G.M. and S.L.; Investigation, G.M., A.T., and S.L.; writing—original draft, G.M. and S.L.; writing—review & editing, G.M. and S.L.; funding acquisition, S.L.; resources, S.L.; supervision, S.L.

## DECLARATION OF INTERESTS

Authors declare that they have no competing interests.

## SUPPLEMENTAL INFORMATION

Supplement 1: Sampling physics in scRNASeq and spRNASeq (theory, supplementary figures S1.1-S1.6, and appendix containing 4 notes with additional statistical/mathematical details and proofs).

Supplement 2: Supplementary figures S2.1, S2.2

## STAR★METHODS

KEY RESOURCES TABLE.

KEY RESOURCES TABLE
| REAGENT OR RESOURCE | SOURCE | IDENTIFIER |
| --- | --- | --- |
| Deposited data |  |  |
| Mouse E18 cortex cells (hippocampus and subventricular zone) | 10X Genomics | See below |
| https://www.10xgenomics.com/datasets/10-k-mouse-e-18-combined-cortex-hippocampus-and-subventricular-zone-cells-single-indexed-3-1-standard-4-0-0 |  |  |
| Mouse E18 cortex cells (hippocampus and subventricular zone) | 10X Genomics | See below |
| https://www.10xgenomics.com/datasets/10-k-mouse-e-18-combined-cortex-hippocampus-and-subventricular-zone-cells-dual-indexed-3-1-standard-4-0-0 |  |  |
| Experimental models: Organisms/strains |  |  |
| C57BL/6J mice maintained on C57BL/6J background | Jackson Laboratory | stock #000664 |
| Critical commercial assays |  |  |
| Visium HD | 10X Genomics | cat #1000676 |
| Software and algorithms |  |  |
| R-Studio | Posit Software | v. 2024.12.0 Build 467 |
| Seurat | Satja Lab<br>(https://satijalab.org/seurat/) | v. 5.2.0 |
| SigmaPlot | Systat Software | v. 13.0 |

## METHOD DETAILS

### Animals

C57BL/6J (stock #000664) mice were purchased from the Jackson Laboratory (Bar Harbor, ME, USA) and maintained on the C57BL/6J background. Animal care and experiments were performed in accordance with a protocol approved by the Eunice Kennedy Shriver National Institute of Child Health and Human Development Animal Care and Use Committee.

### Visium HD experiment

Tibias from mouse embryo at embryonic day E18.5 were dissected, fixed in freshly prepared 4% methanol-free formaldehyde solution in phosphate-buffered saline for 24 h at room temperature, and embedded into paraffin. Three sections from the same tibia were placed on the same slide within 6.5×6.5 mm area, stained with Hematoxylin and Eosin, imaged on a Nikon Ti2-E inverted microscope with a 20X/0.8NA objective and analyzed with the Visium HD™ assay (10X Genomics) as described by the manufacturer. The sequencing statistics was: 356,837,522 reads, 1080 mean reads per 8×8 µm bin, 89.3% valid barcodes, 99.9% valid UMIs, 28.5% sequencing saturation. A low-resolution image of the sequenced area captured by the CytAssist instrument (10X Genomics) was replaced with the high-resolution Nikon Ti2-E image as described by the manufacturer using Loupe Browser software (10X Genomics). The data were processed with SpaceRanger v.3.0.1 software (10X Genomics). Proliferating chondrocyte areas in the proximal growth plates of the 3 tibias were selected in the Loupe Browser software. The barcodes of 8×8 µm bins within these areas were exported from Loupe Browser for further data processing in R (see data analysis).

### Mouse brain scRNASeq datasets

scRNASeq datasets from E18 mouse brain were downloaded from the 10X Genomics (see Key resources table). Filtered count matrices from the two datasets (filtered features .h5 files) were imported into Seurat objects and merged without renormalization to increase the number of neurons in subsequent DGE analysis. GABAergic neurons were selected based on nonzero counts of *Gad1*, *Gad2*, and *Slc6a1*. Glutamatergic neurons were selected based on nonzero counts of *Gls*, *Grin2b*, and *Slc17a7*.

### Simulated random noise and gene upregulation

To test effects of well-defined noise and changes in gene expression, the *N_Z,i_* matrix of glutamatergic neuron gene counts was modified by either random noise [*N_Z,i_* + *f*_n_(α, *N_Z,i_*)] or noise and expression up(down)regulation [*N_Z,i_* + *f*_n_(α, *N_Z,i_*) + *f*_ex_(*Z*, *i*)]. The noise function *f*_n_(α, *N_Z,i_*),

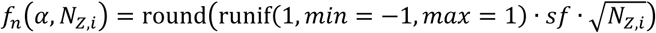

produces integer values uniformly distributed between 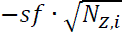 and 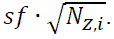. The scaling factor

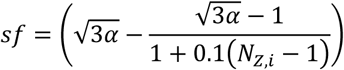

is the simplest expression ensuring 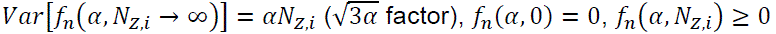 at α < 10, and maximum noise amplitude 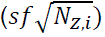 between 0.5*N_Z,i_* and *N_Z,i_* for moderate expression genes with average 0.3 to 3 counts per cell. For the latter genes, simulation results were qualitatively similar at all α < 10. The effects of noise weakly increased with α, and we selected mid-range α = 5 for presenting the results. The results were affected only by the relative noise amplitude but not the specific form of its dependence on *N_Z,i_*.

The expression up(down)regulation function

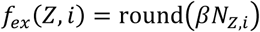

produces integers proportional to the original transcript count *N_Z,i_*, e.g. *β* = 1.5 is used to model 50% upregulation of gene expression.

## QUANTIFICATION AND STATISTICAL ANALYSIS

All data analysis was performed in R-Studio v. 2024.12.0 Build 467 (Posit Software) using R v. 4.4.2, Seurat v. 5.2.0 3 and custom R code shared at https://github.com/sergeyleikin/sc-sp-RNASeq. All statistical tests were performed as described in the main text and Supplement 1. All *t*-tests and weighted *t-*tests were two-tailed with unequal variances. SigmaPlot v.13.0 (Systat Software) was utilized for generating all plots.

## Notes

https://www.ncbi.nlm.nih.gov/geo

https://doi.org/10.7303/syn66520062

https://github.com/sergeyleikin/sc-sp-RNASeq

### Competing Interest Statement

The authors have declared no competing interest.

### Summary of Updates

Minor updates of the manuscript main text based on user and coauthor feedback, which was received and processed after reopening of the US Government. New Appendix in Supplement 1 to provide detailed, step-by-step mathematical derivations and rigorously prove nontrivial formulas (in response to questions posed by users and paper reviewers).

## REFERENCES

1. Squair, J.W., Gautier, M., Kathe, C., Anderson, M.A., James, N.D., Hutson, T.H., Hudelle, R., Qaiser, T., Matson, K.J.E., Barraud, Q., et al. (2021). Confronting false discoveries in single-cell differential expression. Nat Commun 12, 5692. 10.1038/s41467-021-25960-2.

2. Murphy, A.E., Fancy, N., and Skene, N. (2023). Avoiding false discoveries in single-cell RNA-seq by revisiting the first Alzheimer’s disease dataset. Elife 12. 10.7554/eLife.90214.

3. Hao, Y., Stuart, T., Kowalski, M.H., Choudhary, S., Hoffman, P., Hartman, A., Srivastava, A., Molla, G., Madad, S., Fernandez-Granda, C., and Satija, R. (2024). Dictionary learning for integrative, multimodal and scalable single-cell analysis. Nat Biotechnol 42, 293–304. 10.1038/s41587-023-01767-y.

4. Stuart, T., Butler, A., Hoffman, P., Hafemeister, C., Papalexi, E., Mauck, W.M., 3rd, Hao, Y., Stoeckius, M., Smibert, P., and Satija, R. (2019). Comprehensive Integration of Single-Cell Data. Cell 177, 1888–1902 e1821. 10.1016/j.cell.2019.05.031.

5. Chen, Y., Chen, L., Lun, A.T.L., Baldoni, P.L., and Smyth, G.K. (2025). edgeR v4: powerful differential analysis of sequencing data with expanded functionality and improved support for small counts and larger datasets. Nucleic Acids Res 53. 10.1093/nar/gkaf018.

6. Wolf, F.A., Angerer, P., and Theis, F.J. (2018). SCANPY: large-scale single-cell gene expression data analysis. Genome Biol 19, 15. 10.1186/s13059-017-1382-0.

7. Hafemeister, C., and Satija, R. (2019). Normalization and variance stabilization of single-cell RNA-seq data using regularized negative binomial regression. Genome Biol 20, 296. 10.1186/s13059-019-1874-1.

8. Wu, C.H., Zhou, X., and Chen, M. (2025). Exploring and mitigating shortcomings in single-cell differential expression analysis with a new statistical paradigm. Genome Biol 26, 58. 10.1186/s13059-025-03525-6.

9. Tzec-Interián, J.A., González-Padilla, D., and Góngora-Castillo, E.B. (2025). Bioinformatics perspectives on transcriptomics: A comprehensive review of bulk and single-cell RNA sequencing analyses. Quant Biol 13. ARTN e78 10.1002/qub2.78.

10. Gatlin, V., Gupta, S., Romero, S., Chapkin, R.S., and Cai, J.J. (2025). Exploring cell-to-cell variability and functional insights through differentially variable gene analysis. NPJ Syst Biol Appl 11, 29. 10.1038/s41540-025-00507-z.

11. Kim, M.C., Gate, R., Lee, D.S., Tolopko, A., Lu, A., Gordon, E., Shifrut, E., Garcia-Nieto, P.E., Marson, A., Ntranos, V., and Ye, C.J. (2024). Method of moments framework for differential expression analysis of single-cell RNA sequencing data. Cell 187, 6393–6410 e6316. 10.1016/j.cell.2024.09.044.

12. Silkwood, K., Dollinger, E., Gervin, J., Atwood, S., Nie, Q., and Lander, A.D. (2024). Leveraging gene correlations in single cell transcriptomic data. BMC Bioinformatics 25, 305. 10.1186/s12859-024-05926-z.

13. Lee, H., and Han, B. (2024). Pseudobulk with proper offsets has the same statistical properties as generalized linear mixed models in single-cell case-control studies. Bioinformatics 40. 10.1093/bioinformatics/btae498.

14. Yan, J., Zeng, Q., and Wang, X. (2024). RankCompV3: a differential expression analysis algorithm based on relative expression orderings and applications in single-cell RNA transcriptomics. BMC Bioinformatics 25, 259. 10.1186/s12859-024-05889-1.

15. Missarova, A., Dann, E., Rosen, L., Satija, R., and Marioni, J. (2024). Leveraging neighborhood representations of single-cell data to achieve sensitive DE testing with miloDE. Genome Biol 25, 189. 10.1186/s13059-024-03334-3.

16. Han, G., Yan, D., Sun, Z., Fang, J., Chang, X., Wilson, L., and Liu, Y. (2024). Bayesian-frequentist hybrid inference framework for single cell RNA-seq analyses. Hum Genomics 18, 69. 10.1186/s40246-024-00638-0.

17. Ospina, O.E., Soupir, A.C., Manjarres-Betancur, R., Gonzalez-Calderon, G., Yu, X., and Fridley, B.L. (2024). Differential gene expression analysis of spatial transcriptomic experiments using spatial mixed models. Sci Rep 14, 10967. 10.1038/s41598-024-61758-0.

18. Mason, K., Sathe, A., Hess, P.R., Rong, J., Wu, C.Y., Furth, E., Susztak, K., Levinsohn, J., Ji, H.P., and Zhang, N. (2024). Niche-DE: niche-differential gene expression analysis in spatial transcriptomics data identifies context-dependent cell-cell interactions. Genome Biol 25, 14. 10.1186/s13059-023-03159-6.

19. Cuevas-Diaz Duran, R., Wei, H., and Wu, J. (2024). Data normalization for addressing the challenges in the analysis of single-cell transcriptomic datasets. BMC Genomics 25, 444. 10.1186/s12864-024-10364-5.

20. Pullin, J.M., and McCarthy, D.J. (2024). A comparison of marker gene selection methods for single-cell RNA sequencing data. Genome Biol 25, 56. 10.1186/s13059-024-03183-0.

21. Liu, Y., Zhao, J., Adams, T.S., Wang, N., Schupp, J.C., Wu, W., McDonough, J.E., Chupp, G.L., Kaminski, N., Wang, Z., and Yan, X. (2023). iDESC: identifying differential expression in single-cell RNA sequencing data with multiple subjects. BMC Bioinformatics 24, 318. 10.1186/s12859-023-05432-8.

22. Cui, T., and Wang, T. (2023). A comprehensive assessment of hurdle and zero-inflated models for single cell RNA-sequencing analysis. Brief Bioinform 24. 10.1093/bib/bbad272.

23. Liu, Y., Huang, J., Pandey, R., Liu, P., Therani, B., Qiu, Q., Rao, S., Geurts, A.M., Cowley, A.W., Jr., Greene, A.S., and Liang, M. (2023). Robustness of single-cell RNA-seq for identifying differentially expressed genes. BMC Genomics 24, 371. 10.1186/s12864-023-09487-y.

24. Gagnon, J., Pi, L., Ryals, M., Wan, Q., Hu, W., Ouyang, Z., Zhang, B., and Li, K. (2022). Recommendations of scRNA-seq Differential Gene Expression Analysis Based on Comprehensive Benchmarking. Life (Basel) 12. 10.3390/life12060850.

25. Shi, Y., Lee, J.H., Kang, H., and Jiang, H. (2022). A Two-Part Mixed Model for Differential Expression Analysis in Single-Cell High-Throughput Gene Expression Data. Genes (Basel) 13. 10.3390/genes13020377.

26. Bouland, G.A., Mahfouz, A., and Reinders, M.J.T. (2021). Differential analysis of binarized single-cell RNA sequencing data captures biological variation. NAR Genom Bioinform 3, lqab118. 10.1093/nargab/lqab118.

27. Li, H., Zhu, B., Xu, Z., Adams, T., Kaminski, N., and Zhao, H. (2021). A Markov random field model for network-based differential expression analysis of single-cell RNA-seq data. BMC Bioinformatics 22, 524. 10.1186/s12859-021-04412-0.

28. Fleiss, J.L. (1999). The Design and Analysis of Clinical Experiments (John Wiley & Sons).

29. Hayes, R.J., and Moulton, L.H. (2022). Cluster Randomized Trials (CRC Press).

30. Ridout, M.S., Demetrio, C.G., and Firth, D. (1999). Estimating intraclass correlation for binary data. Biometrics 55, 137–148. 10.1111/j.0006-341x.1999.00137.x.

31. Hartung, J., Knapp, G., and Sinha, B. (2008). Statistical Meta-Analysis with Applications (Wiley-Interscience).

32. Bevington, P.R., and Robinson, D.K. (2002). Data Reduction and Error Analysis for the Physical Sciences, Third Edition Edition (McGraw-Hill).

33. Ramachandran, K.M., and Tsokos, C.P. (2021). Mathematical Statistics with Applications in R, Third Edition Edition (Academic Press).

34. Jiang, R., Sun, T., Song, D., and Li, J.J. (2022). Statistics or biology: the zero-inflation controversy about scRNA-seq data. Genome Biol 23, 31. 10.1186/s13059-022-02601-5.

35. Silverman, J.D., Roche, K., Mukherjee, S., and David, L.A. (2020). Naught all zeros in sequence count data are the same. Comput Struct Biotechnol J 18, 2789–2798. 10.1016/j.csbj.2020.09.014.

36. Zhao, P., Zhu, J., Ma, Y., and Zhou, X. (2022). Modeling zero inflation is not necessary for spatial transcriptomics. Genome Biol 23, 118. 10.1186/s13059-022-02684-0.

37. Svensson, V. (2020). Droplet scRNA-seq is not zero-inflated. Nat Biotechnol 38, 147–150. 10.1038/s41587-019-0379-5.

38. Lun, A.T.L., and Marioni, J.C. (2017). Overcoming confounding plate effects in differential expression analyses of single-cell RNA-seq data. Biostatistics 18, 451–464. 10.1093/biostatistics/kxw055.

39. Love, M.I., Huber, W., and Anders, S. (2014). Moderated estimation of fold change and dispersion for RNA-seq data with DESeq2. Genome Biol 15, 550. 10.1186/s13059-014-0550-8.

40. Rasch, D., Teuscher, F., and Guiard, V. (2007). How robust are tests for two independent samples? J Stat Plan Infer 137, 2706–2720. 10.1016/j.jspi.2006.04.011.

41. Hoffman, G.E., Lee, D., Bendl, J., Prashant, N.M., Hong, A., Casey, C., Alvia, M., Shao, Z., Argyriou, S., Therrien, K., et al. (2024). Efficient differential expression analysis of large-scale single cell transcriptomics data using dreamlet. bioRxiv. 10.1101/2023.03.17.533005.

42. Hoffman, G.E., and Roussos, P. (2021). Dream: powerful differential expression analysis for repeated measures designs. Bioinformatics 37, 192–201. 10.1093/bioinformatics/btaa687.

43. Amezquita, R.A., Lun, A.T.L., Becht, E., Carey, V.J., Carpp, L.N., Geistlinger, L., Marini, F., Rue-Albrecht, K., Risso, D., Soneson, C., et al. (2020). Orchestrating single-cell analysis with Bioconductor. Nat Methods 17, 137–145. 10.1038/s41592-019-0654-x.

44. Benjamini, Y., and Hochberg, Y. (1995). Controlling the False Discovery Rate – a Practical and Powerful Approach to Multiple Testing. J R Stat Soc B 57, 289–300. DOI 10.1111/j.2517-6161.1995.tb02031.x.

45. Benjamini, Y., and Cohen, R. (2017). Weighted false discovery rate controlling procedures for clinical trials. Biostatistics 18, 91–104. 10.1093/biostatistics/kxw030.

