## Supplement 1 for "Differential expression analysis in single cell and spatial RNASeq without model assumptions"

### Sampling physics in scRNASeq and spRNASeq

The scRNASeq data matrix $N_{Z,i}$ of gene $Z$ counts in cell $i$ is widely interpreted assuming transcript sampling probability being consistent from cell to cell in the original biological sample, e.g., for data parametrization within generalized linear models (GLMs) of count statistics [1-5]. However, these are desired rather than actual data properties. The physics of sampling, technical variability (hereafter noise), and biological variability of cells suggest that the statistics of $N_{Z,i}$ cannot be described within GLMs or other simplified parametric models. In this supplement, we explain why this is the case and derive nonparametric statistics for calculating average expression, its variance, and probability of differential expression. At the end, we discuss distinct features of spRNASeq assays, and the corresponding modifications needed for their analysis.

##### Sampling and technical noise

Consider common droplet-based scRNASeq assays. All of them involve the following basic steps: *Step 1.* Cell extraction from tissue and dissociation. *Step 2*. Encapsulation of cells inside droplets containing primers and reagents, one cell per droplet. *Step 3.* Cells lysis, RNA hybridization with primers, and synthesis of barcoded cDNAs by reverse transcription. A primer-based barcode includes a sequence identifying the droplet and cell inside as well as a sequence unique for each primer (unique molecular identifier, UMI). *Step 4.* Purification and PCR amplification of cDNAs extracted from droplets. *Step 5.* Counting of transcripts in an aliquot of the resulting cDNA solution (cDNA library) by next generation sequencing (NGS). Each cDNA is mapped to a gene based on the transcript sequence and to an individual cell based on the droplet barcode. Each unique transcript is counted only once based on its UMI.

The NGS counting can indeed be described by traditional counting statistics, since the probability of each cDNA being counted is approximately the same. However, being remarkably efficient and more than 90% accurate [6], it is not the main source of technical noise in scRNASeq. The only logical explanation of the variation from ~ $10^3$ up to ~ $10^5$ RNA counts per cell (Fig. 1A) is that the preceding assay steps produce much larger technical noise. Among these technical noise sources is initial cell isolation and encapsulation into droplets, which alters the composition of transcripts inside cells because cells react to enzymes, disruption of cell-cell contacts, etc. These transcript composition changes are expected to result in significant cell-to-cell variations in the sampling probability for the same gene. These variations are further increased by fluctuations in primer and reagent composition and concentration from droplet to droplet, variable RNA degradation, variable likelihood of hybridization with primers, and variable reverse transcription efficiency. All these noise sources and biological cell variability add up to the observed ~ 100-fold variation in the transcript sampling probability from cell to cell, which is not likely to be amenable to parametrization within GLMs or other simplified models.

As we show below, this complex physics is better described by calculating the average of relative counts $n_{Z,i} = N_{Z,i}/\sum_Z N_{Z,i}$ and its variance, without any oversimplification and parametrization. We argue that the only assumption needed for determining $\log_2(\mathrm{FC})$ and the corresponding $p$-values in DGE analysis is that all transcript counts are random and independent (as also assumed in other statistical analyses of scRNASeq). Such random, independent sampling of transcripts relies on low fraction of all transcripts in the cell being detected by the current technology. Indeed, a large fraction of live cell transcripts is expected to be degraded/damaged during cell encapsulation into droplets and lysis. Only some of the surviving transcripts are then converted into high quality (identifiable) cDNA, and usually no more than ~50% of the latter are sequenced by NGS at the recommended sequencing depth. When all these effects are combined, the upper estimate for the sampling fraction in scRNASeq is ~ 1-10%, justifying the assumption of random and independent sampling. Consistently, from $10^4$ to $3\cdot10^4$ unique transcripts per cell are sequenced in most cells in Fig. 1A. A mammalian cell is estimated to have ~ $10^5$ – $10^6$ transcripts depending on its transcriptional activity [7,8]. GABAergic and glutamatergic neurons are large, highly transcriptionally active cells likely to contain no less than ~ $10^6$ transcripts.

All scRNASeq analysis methods assume random, independent transcript sampling. Unlike other methods, however, our approach does not utilize additional assumptions. Instead, we rely on the analysis framework developed for studies of cluster-randomized experiments [9-11], since each cell is analogous to a cluster of independent transcript sequencing experiments. We adapt and modify this approach to account for the specific physical properties of scRNASeq assays.

**Average and variance calculation**

To calculate the average and variance, we use that Eq. (1) for $n_{Z,i}$ can be rewritten in the following form

$$n_{Z,i} = \frac{1}{N_i}\sum_{j=1}^{N_i} X_{Z,i,j} \qquad (S1)$$

where $X_{Z,i,j} = 0{,}1$ is gene $Z$ count from cell/droplet $i$ and UMI $j$, $n_{Z,i}$ is the total normalized count of gene $Z$ in cell $i$ and $N_i = \sum_Z N_{Z,i}$ is the total number of unique transcripts (UMIs) detected in cell $i$ (for discussion of failed transcript identifications see Appendix, Note 1).

One may be tempted to interpret $X_{Z,i,j}$ in terms of common counting models, e.g., as a Bernoulli trial, yet this may not be consistent with scRNASeq realities, and this is not what we do. Such models do not account for continuous and rapid change in the number and composition of transcripts, which happens both in a live cell and when the transcripts are being sampled after the cell is lysed. Moreover, even for given total number of transcripts in the cell ($N_i^{tot}$), total number of gene $Z$ transcripts ($N_{Z,i}^{tot}$), and sampling failure rate ($FR_i$, fraction of transcripts destroyed during sampling but not identified), the probability density for $N_{Z,i}$ may not be just a function of $N_i$, $N_i^{tot}$, $N_{Z,i}^{tot}$, and $FR_i$ (as assumed in all counting models). Instead, it is also likely to depend on transcript distribution in different locations (e.g., nucleus, cytosol, ribosomes), their physical state (e.g., folding, splicing, and interactions), concentration and composition of sequencing reagents, etc. In contrast, a textbook model of sampling of colored balls from a basket either has a constant outcome probability (canonical Bernoulli trials of sampling with replacement) or a probability determined only by the outcome ($N_{Z,i}$ and $N_i$), and total numbers of balls ($N_{Z,i}^{tot}$, $N_i^{tot}$), which is unaffected by the spatial distribution of balls in the basket. We therefore use the approach we think is better justified, which is to assume that the sampling events are random and independent without making any other assumptions. Then, our results are not limited just to the common counting models.

Very large variation in $N_i$ from cell to cell (Fig. 1A) indicates large variation in the precision of measuring $n_{Z,i}$ due to the technical noise, in which case the average value of $n_{Z,i}$ in a group of $N$ cells must be calculated as a weighted average [9-12]

$$\bar{n}_Z = \sum_{i=1}^{N} n_{Z,i} w_{Z,i} \;, \tag{S2}$$

where

$$w_{Z,i} = \frac{1}{Var(n_{Z,i})} \Big/ \sum_{i=1}^{N} \frac{1}{Var(n_{Z,i})} \tag{S3}$$

is the statistical weight of gene $Z$ transcripts in cell $i$ with $N_i$ transcripts ($\sum_i w_{Z,i} = 1$). $Var(n_{Z,i})$ is the total variance of $n_{Z,i}$,

$$Var(n_{Z,i}) = \langle \left(n_{Z,i} - E(n_Z|N_i)\right)^2 \rangle \tag{S4}$$

which must be calculated by averaging over all possible statistical trajectories producing a cell with $N_i$ transcripts. Here $E(n_Z|N_i)$ is the expected value of $n_Z$ in any cell of the selected type that has $N_i$ transcripts. The symbol $\langle \quad \rangle$ indicates statistical trajectory averaging (akin to Feynman's path integrals in statistical physics [13]).

Using the law of total variance [14], we can calculate the variance of $n_{Z,i}$ as

$$Var(n_{Z,i}) = \langle \left(n_{Z,i} - \langle n_{Z,i} \rangle_i\right)^2 \rangle + V_Z \tag{S5}$$

where $\langle n_{Z,i} \rangle_i$ is the expected value of $n_{Z,i}$ (average over statistical trajectories in the cell *i*) and $V_Z = \langle \left(\langle n_{Z,i} \rangle_i - E(n_Z|N_i)\right)^2 \rangle$. After substituting Eq. ($S1$) into Eq. ($S5$) and calculating the sums, we find

$$Var(n_{Z,i}) = \langle \frac{\langle n_{Z,i} \rangle_i \left(1 - \langle n_{Z,i} \rangle_i\right)}{N_i} \rangle + V_Z = \frac{\bar{n}_Z(1 - \bar{n}_Z) + V_Z(N_i - 1)}{N_i} \tag{S6}$$

where we used that $Var(x) = \langle x^2 \rangle - \langle x \rangle^2$ and $\bar{n}_Z \cong \langle n_{Z,i} \rangle$. Substituting Eq. ($S6$) into ($S3$), we find

$$w_{Z,i} = \frac{N_i}{\bar{n}_Z(1 - \bar{n}_Z) + (N_i - 1)V_Z} \Big/ \sum_{i=1}^{N} \frac{N_i}{\bar{n}_Z(1 - \bar{n}_Z) + (N_i - 1)V_Z} \,. \tag{S7}$$

Note that we derived Eq. (*S*6) by directly calculating the sums, assuming only that all sequencing counts $X_{Z,i,j}$ are random and independent (for detailed calculation of the sums see Appendix, Note 2). We did not treat the sequencing reads as Bernoulli trials or utilize any specific statistical distribution. Another instructive way to arrive at Eq. (*S*6) would be to treat $X_{Z,i,j}$ as a count of gene $Z$ in a random draw $j$ from the total population of $N_i^{tot}$ transcripts in the cell *i*. Then $\sum_j X_{Z,i,j}$ is described by the hypergeometric distribution, in which $N_i$ is the total number of random draws (conversion to cDNA, sequencing, and identification of a unique transcript). Even though the value of $N_i^{tot}$ is not known, $Var(n_{Z,i})$ is then described by Eq. (*S*6) as long as $N_i \ll N_i^{tot}$, which is the case in typical scRNASeq experiments (Appendix, Notes 3 and 4).

Eq. (*S*7) is identical to

$$w_{Z,i} = \frac{N_i}{1 + \rho_Z(N_i - 1)} \Big/ \sum_{i=1}^{N} \frac{N_i}{1 + \rho_Z(N_i - 1)} \,, \tag{S8}$$

where $\rho_Z = V_Z/\bar{n}_Z(1-\bar{n}_Z)$ is the intracluster correlation coefficient ICC ($0 \le \rho_Z \le 1$). Eq. ($S8$) is well-known in statistics of cluster-randomized experiments [9,11,15-17]. Previous models of scRNASeq data analysis correspond to $\rho_Z = 1$ ($w_{Z,i} = 1/N$) and $\rho_Z = 0$ ($w_{Z,i} = N_i/\sum_i N_i$). Statistical studies of cluster-randomized experiments showed that neither of these two limits is a good approximation. The importance of accurate ICC calculation has been recognized, and multiple different ICC estimator models have been developed [17]. The most widely used is the ANOVA ICC estimator [15-17]. Here we suggest a different approach for estimating $\rho_Z$ directly from experimental data without relying on any specific type of data distribution or sampling. The benefit of this approach is that it makes no assumptions and can also be used for spRNASeq and multiple sample analysis in scRNASeq.

To calculate $\rho_Z$ we use that for inverse variance weights [10,12]

$$Var(\bar{n}_Z) = 1\Big/\sum_{i=1}^{N}\left[1/Var(n_{Z,i})\right] = \bar{n}_Z(1-\bar{n}_Z)\Big/\sum_{i=1}^{N}\left(\frac{N_i}{1+\rho_Z(N_i-1)}\right). \qquad (S9)$$

At the same time, $Var(\bar{n}_Z)$ can be estimated directly from the data for given inverse-variance weights $w_{Z,i}$ using the unbiased estimator

$$Var(\bar{n}_Z) = \frac{1}{1-\sum_{i=1}^{N} w_{Z,i}^2}\sum_{i=1}^{N} w_{Z,i}^2\left(n_{Z,i}-\bar{n}_Z\right)^2. \qquad (S10)$$

Taken together, Eqs. ($S8$)-($S10$) allow calculation of $w_{Z,i}$ directly from the data without any assumptions as we show below.

While Eqs. ($S8$) and ($S9$) are well known in the theory of cluster-randomized experiments, we have not seen Eq. ($S10$) being used before. To derive this variance estimator, we used that

$$\langle\sum_{i=1}^{N} w_{Z,i}^2\left(n_{Z,i}-\bar{n}_Z\right)^2\rangle = \sum_{i=1}^{N} w_{Z,i}^2\langle n_{Z,i}^2\rangle + \sum_{i=1}^{N} w_{Z,i}^2\langle \bar{n}_Z^2\rangle - 2\sum_{i=1}^{N} w_{Z,i}^2\langle n_{Z,i}\bar{n}_Z\rangle \qquad (S11)$$

It follows from Eqs. ($S7$),($S9$) that

$$w_{Z,i}Var(n_{Z,i}) = Var(\bar{n}_Z) \qquad (S12)$$

Therefore,

$$w_{Z,i}^2\langle n_{Z,i}^2\rangle = w_{Z,i}^2 Var(n_{Z,i}) + w_{Z,i}^2\langle \bar{n}_Z\rangle^2 = w_{Z,i}Var(\bar{n}_Z) + w_{Z,i}^2\langle \bar{n}_Z\rangle^2, \qquad (S13)$$

$$w_{Z,i}^2\langle \bar{n}_Z^2\rangle = w_{Z,i}^2 Var(\bar{n}_Z) + w_{Z,i}^2\langle \bar{n}_Z\rangle^2, \qquad (S14)$$

and

$$w_{Z,i}^2\langle n_{Z,i}\bar{n}_Z\rangle = w_{Z,i}^3 Var(n_{Z,i}) + w_{Z,i}^2\langle \bar{n}_Z\rangle^2 = w_{Z,i}^2 Var(\bar{n}_Z) + w_{Z,i}^2\langle \bar{n}_Z\rangle^2. \qquad (S15)$$

After substitution of Eqs. (S12)-($S15$) into Eq. ($S11$) we arrive at

$$Var(\bar{n}_Z) = \frac{1}{\left(1-\sum_{i=1}^{N} w_{Z,i}^2\right)}\langle\sum_{i=1}^{N} w_{Z,i}^2\left(n_{Z,i}-\bar{n}_Z\right)^2\rangle, \qquad (S16)$$

which is approximated by Eq. ($S10$) in the limit of large $N$.

It is worth noting that Eqs. ($S10$) and ($S16$) are approximations when the weights $w_{Z,i}$ are fitted from the data rather than fixed. However, this is justified when uncertainties of $w_{Z,i}$ are much smaller than variations of $n_{Z,i}$, which should be the case for $w_{Z,i}$ calculated based on averaging over multiple cells as described below.

Combining Eqs. ($S9$) and ($S10$) we arrive at

$$\bar{n}_Z(1-\bar{n}_Z)\Big/\sum_{i=1}^{N}\left(\frac{N_i}{1+\rho_Z(N_i-1)}\right)=\frac{1}{1-\sum_{i=1}^{N}w_{Z,i}^2}\sum_{i=1}^{N}w_{Z,i}^2\left(n_{Z,i}-\bar{n}_Z\right)^2. \qquad (S17)$$

Using Eq. ($S8$) to express $w_{Z,i}$ as a function of $\rho_Z$ and $N_i$ and Eq. ($S2$) to express $\bar{n}_Z$ as a function of $w_{Z,i}$ and $n_{Z,i}$, Eq. ($S17$) can be solved numerically to find $\rho_Z$. The values of $\bar{n}_Z$ and $Var(\bar{n}_Z)$ calculated from Eq. ($S17$) are in good agreement with those calculated by using the ANOVA ICC estimator (Fig. $S1.1$). Because the latter calculation is computationally faster, we provide it as an option in our scRNASeq data analysis functions (see https://github.com/sergeyleikin/sc-sp-RNASeq). Note that Eq. ($S17$) may yield negative values of $\rho_Z$ instead of $\rho_Z \approx 0$ (like ANOVA ICC estimator), which is a known effect of sampling errors that should be corrected by changing negative $\rho_Z$ to $\rho_Z = 0$ [9]. Similarly, estimated $\rho_Z > 1$ resulting from sampling errors should be replaced with $\rho_Z = 1$ (although we have not observed estimated $\rho_Z > 1$ in our analysis of experimental data or simulations).

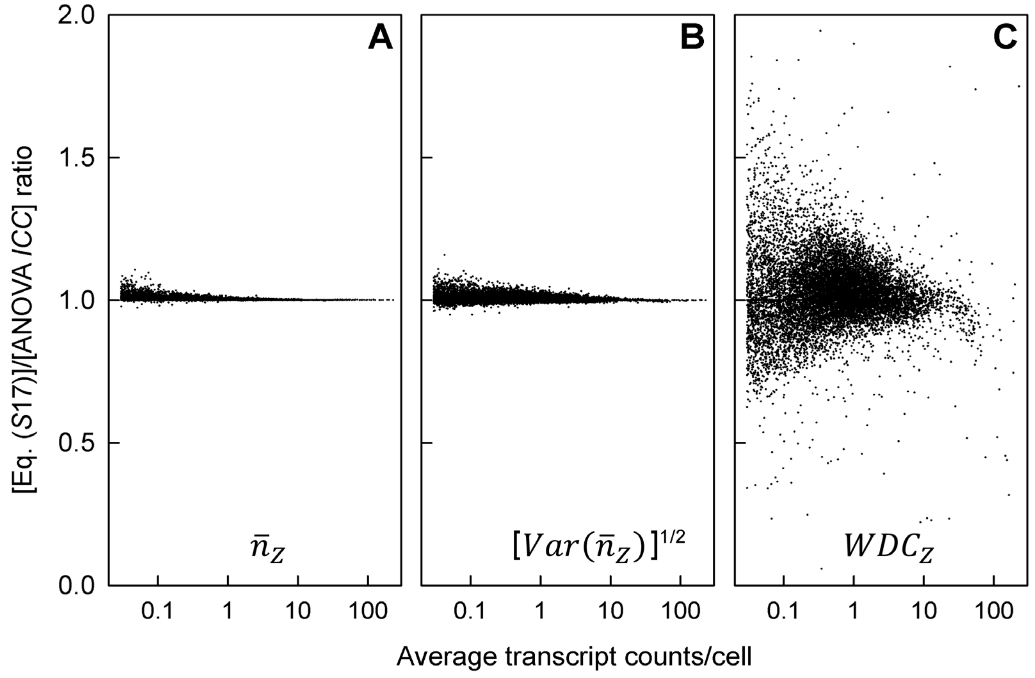


**Figure S1.1. Calculation of $\bar{n}_Z$, $[Var(\bar{n}_Z)]^{1/2}$, and statistical weight distribution coefficient ($WDC_Z$) from Eq. ($S17$) vs. ANOVA ICC.** Ratios of the corresponding values within the glutamatergic neuron dataset (Fig. 1) are plotted vs. $\left(\sum_{i=1}^{N} N_{Z,i}\right)/N$ for each gene $Z$ (data points) with $\left(\sum_{i=1}^{N} N_{Z,i}\right)/N > 0.03$.

Finding $\bar{n}_Z$ and $Var(\bar{n}_Z)$ through data-driven estimates of $w_{Z,i}$ enables calculation of $\log_2(\mathrm{FC})$ and the corresponding $p$-values with increased accuracy and power. The data analysis reported in the main text of the paper and built into our R scripts is based on this approach. Its only underlying assumptions are that the transcript sampling in scRNASeq is random and independent, and that fluctuations in $\langle n_{Z,i}\rangle$ and $N_i$ across cells are independent of each other, e.g., because of large technical noise. While technical noise might not completely remove correlation between $\langle n_{Z,i}\rangle$ and $N_i$ across cells of different types, here we are mostly concerned with DGE between groups of similar cells. These two assumptions are essential for all unbiased models of scRNASeq analysis anyway. The benefit of our method is that unlike the commonly used approaches it does not use any additional assumptions, simplifications, or data parametrization, while taking full advantage of the data at hand.

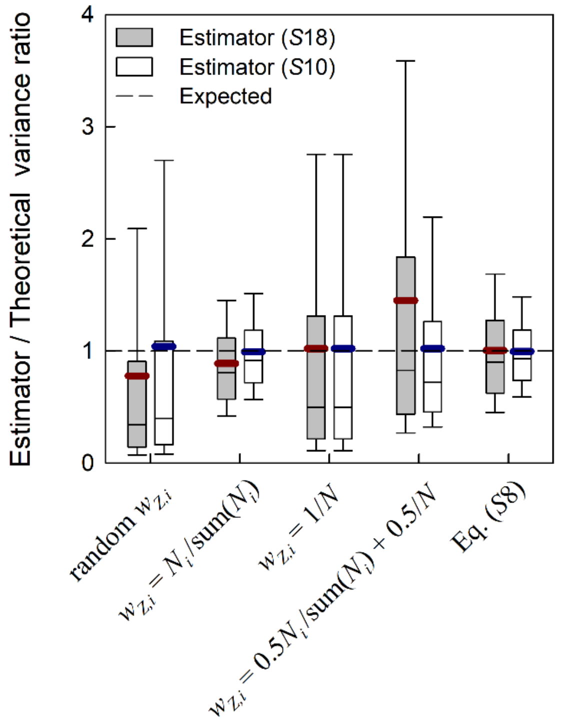


**Figure S1.2. Performance of different variance estimators for weighted average.** Ratios of $Var(\bar{n}_Z)$ estimated from Eqs. ($S$10), ($S$18) to the theoretically expected $Var(\bar{n}_Z) = \sum_{i=1}^{N} w_{Z,i}^2 \left[\langle p_Z\rangle(1-\langle p_Z\rangle) + (N_i - 1)Var(p_Z)\right]/N_i$, where $p_Z$ is the probability distribution function used to simulate $N_{Z,i}$. The simulations consisted of $N_i$ Bernoulli trails within each of $N$=100 data clusters $i$ (proxy for cells) with the beta probability distribution function $p_Z$ that had shape parameters $\alpha$=2 and $\beta$=20,000 to calculate $N_{Z,i}$. To generate random $N_i$ in the range from 100 to 10,000, we utilized $N_i = \mathrm{round}([n_{max}]^r)$ with $n_{max} = 10{,}000$ and $r$ a uniformly distributed random number between 0.5 and 1. The box plots show median, 10$^{th}$, 25$^{th}$, 75$^{th}$, and 90$^{th}$ percentile for 10,000 simulations with each of the indicated $w_{Z,i}$. The bold red and blue bars show mean values for the corresponding estimators.

**Weighted *t*-test**

A variety of tests for comparing weighted average values in cluster-randomized experiments have been described, partly because of suboptimal weighted *t*-test performance when statistical weights are imprecise [9-11]. We have found, however, that weighted *t*-test performance issues are caused by common usage of the following variance estimator

$$Var(\bar{x}) = \frac{1}{N-1}\sum_{i=1}^{N} w_{x,i}(x_i - \bar{x})^2 \quad . \tag{S18}$$

for random variables $x_i$ with weights $w_{x,i}$ in experiment clusters *i*. This expression provides unbiased estimate of $Var(\bar{x})$ at $w_{x,i} = [Var(x_i)]^{-1}/\sum_i [Var(x_i)]^{-1}$ [12]. However, it may become biased and inaccurately estimate $Var(\bar{n}_Z)$ when $w_{Z,i}$ do not match the inverse variance of $n_{Z,i}$ (Fig. S1.2), e.g., resulting in increased false positive findings at $\rho_Z = 0$ (Fig.S1.3A,B,E). Moreover, the test with variance estimator (*S*18) can be anti-conservative even with inverse variance weights (Fig S1.3A,B,E). Apparently, (*S*18) can have larger uncertainty than (*S*10) and in some cases underestimate the variance, causing small *p*-values. We are not aware of any rigorous justification or derivation of the estimator (*S*18).

In contrast, the variance estimator given by Eq. (*S*10) remains unbiased and accurate at any $w_{Z,i}$ provided that $\sum_i w_{Z,i}^2 \ll 1$, which is almost always true for scRNASeq and spRNASeq. After replacing the variance estimator (*S*18) with (*S*10), we found that the weighted *t*-test exhibited optimal or close to optimal type I error (false positive findings) rate control (Fig. S1.3). Since we are aware only of weighted *t*-test implementations based on Eq. (*S*18) we have included an R code for weighted *t*-test based on Eq. (*S*10) variance estimator into our package of R functions for scRNASeq and spRNASeq analysis. In addition, this code has two options for calculating degrees of freedom, which may affect *p*-values in some cases (for more details, see https://github.com/sergeyleikin/sc-sp-RNASeq). It is, however, important to keep in mind that the weighted *t*-test is exact only in the asymptotic limit of large *N*, i.e., at large number of degrees of freedom. Otherwise, it is just a reasonable approximation.

Analysis of experimental data (Figs. 3,4) showed that the weighted *t*-test with the (*S*10) variance estimator might provide the best combination of type I error rate control and type II error (false negative) rate control. However, the ground truth in experimental data is not known. Therefore, we performed extensive comparison of all widely used statistical tests for simulated data, in which the ground truth is known (Figs. S1.3 and S1.4). We simulated single gene $N_{Z,i}$ counts in scRNASeq by using beta distribution for the count probability, varying the first shape parameter α to mimic changes in gene expression. We used a scaling factor $n_{max}$ in the distribution function for the total counts per cell to mimic changes in the sequencing depth. We tested different versions of this distribution, all of which produced similar results (see additional data at https://doi.org/10.7303/syn66520062).

These simulations confirmed our conclusions based on experimental data analysis. The only two tests with good type I error rate control under all conditions were weighted *t*-test with the variance estimator (*S*10) and unweighted (common) *t*-test, although the unweighted *t*-test was too conservative under some conditions (Figs. S1.3A-F, S1.4E-H). The weighted *t*-test with variance estimator (*S*18) had too many false positive findings (Figs. S1.3A,B,E). Consistent with our analysis of real experimental data in scRNASeq, the Wilcoxon test exhibited poor type I error control for samples with different distributions of $N_i$, e.g., mimicking effects of sequencing depth variation (Figs. S1.3A,C,G, S1.4F,H).

The weighted $t$-test with the variance estimator ($S10$) was also among the best performers in detecting true changes in expression, outperforming the Wilcoxon test (Fig. S1.4A,B,D) despite the type I error rate control issues in the latter (Figs. S1.3A,C, S1.4F,H), depending on the underlying distributions. The unweighted $t$-test was too conservative and exhibited the worst type II error rate control among all tests.

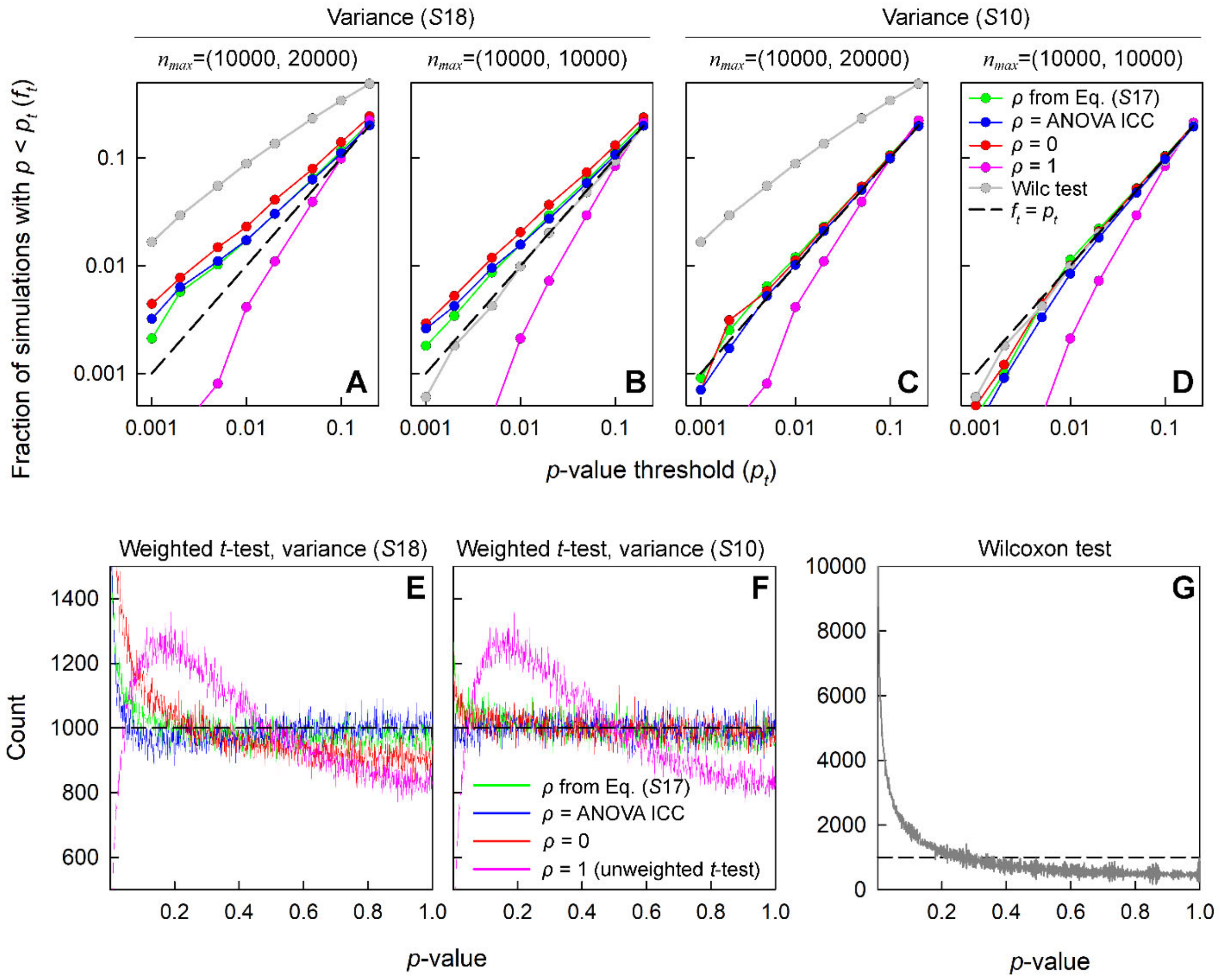


**Figure S1.3. Type I error (false positive findings) in weighted *t*-test, unweighted *t*-test ($\rho = 1$) and Wilcoxon test (Wilc). A-D.** Fractions $f_t$ of 10,000 simulated dataset pairs with $p$-values below $p_t$. The datasets were simulated using the same probability distribution functions and random generator of $N_i$ as in Fig. S1.2. The plots show comparisons of 10,000 dataset pairs that had $\alpha = 2$, $\beta = 20{,}000$, and either different (**A,C**) or the same (**B,D**) $n_{max}$ within each pair. The numbers in parenthesis show the two values of $n_{max}$ within each pair. Variance estimators ($S18$) and ($S10$) were used for the weighted $t$-test as indicated. Tests with $f_t > p_t$ have poor type I error rate control. Tests with $f_t < p_t$ are conservative. **E and F.** Histograms of weighted $t$-test $p$-values comparing 1,000,000 dataset pairs simulated using the same parameters as in A and C, respectively**.** Colored lines show numbers of counts per bin (bin size = 0.001). **G.** Histogram of Wilcoxon test $p$-values in simulations shown in **E** and **F**. Black dashed lines in all panels show expected histograms at optimal type I error rate control.

Overall, the weighted $t$-test with the variance estimator ($S10$) was the clear winner, like in the experimental data analysis (Fig. S1.4). It is worth noting that computationally faster weighted $t$-test with

ANOVA ICC performed nearly identical to the one based on numerical calculation of $\rho_Z$ from Eq. ($S$17). The ANOVA ICC estimator is therefore available as an option in our scRNASeq data analysis functions.

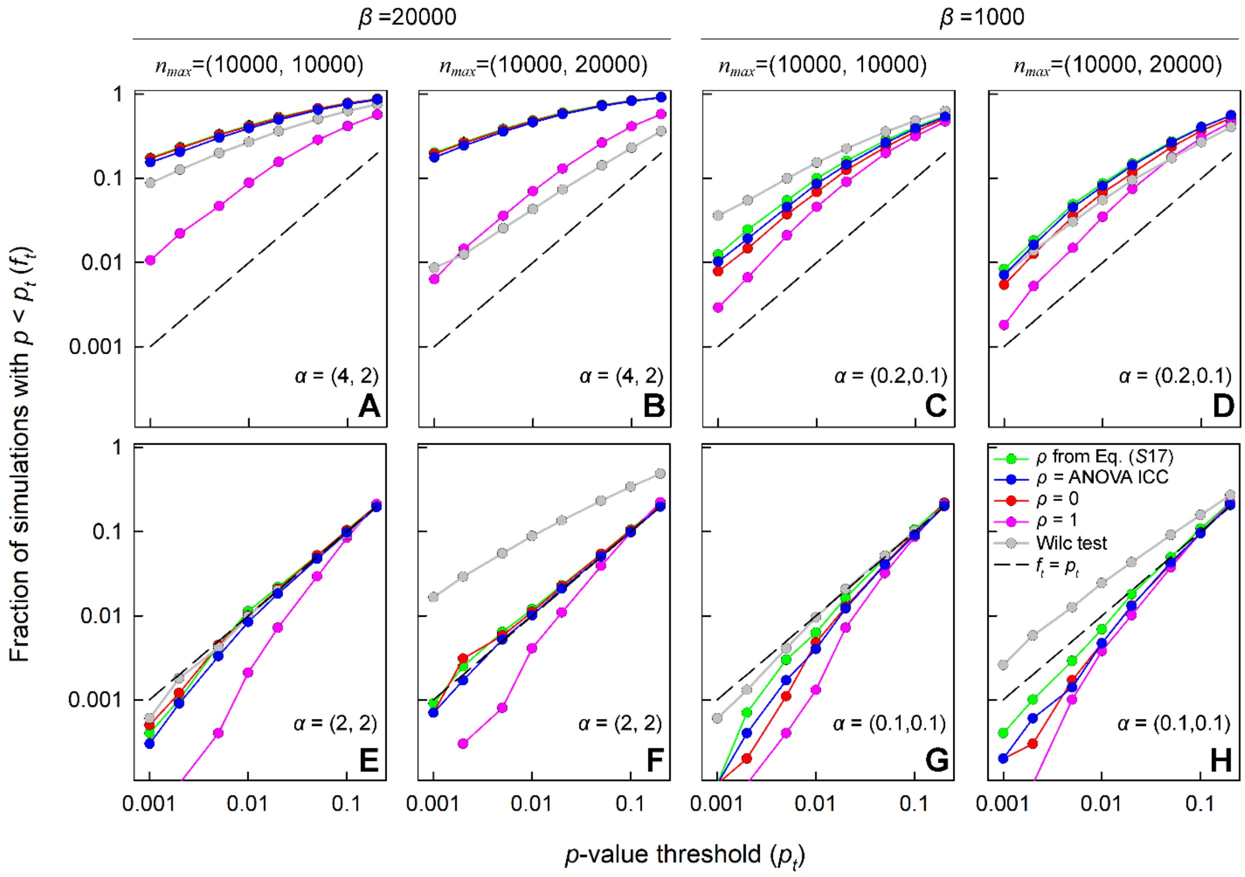


**Figure S1.4. Power to identify true positive findings (top) and false positive rate control (bottom) of weighted *t*-test, unweighted *t*-test ($\rho$ = 1) and Wilcoxon test (Wilc).** For these simulations, we used the same probability distribution functions as in Figs. S1.2 and S1.3 and $N$ = 100. All other parameters were as indicated. Parameter values in parentheses are for the first and second dataset in each of 10,000 simulated dataset pairs. Dashed black lines show $f_t = p_t$ expected at optimal type I error rate control for plots with the same values of $\alpha$ within each dataset pair (bottom panels). Higher fractions $f_t$ of 10,000 simulated dataset pairs with $p$-values below $p_t$ (curves above the dashed lines) indicate increased test power in datasets with different $\alpha$ within each pair (top panels).

#### Violin plot interpretation

Using equal statistical weights of different cells ($\rho_Z$ = 1) may not only distort the calculated $p$-values but also cause misinterpretation of the popular violin plots for normalized gene expression (Fig. S1.5). A violin plot is a vertical histogram of cell density distribution vs. normalized gene expression, which is meaningful only when all cells can be assumed to have the same statistical weight. A histogram peak showing only low confidence cells (with low statistical weights) is not an accurate representation of actual gene expression and may be misleading.

To demonstrate misleading violin plots, Fig. *S*1.5 shows an effect of 40% subsampling of all genes (every count retained with 40% probability or discarded with 60% probability), which corresponds to reduced depth of sequencing in experiments and does not alter gene expression. However, this subsampling significantly alters the violin plots.

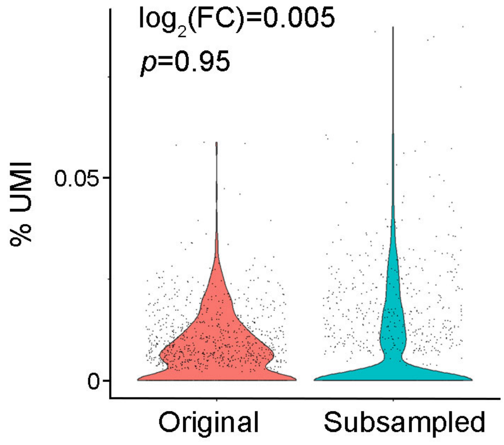


**Figure S1.5. Example of misleading violin plots.** Violin plots for normalized transcript counts of *Taz* in original glutamatergic neuron data (red) and the same data after 40% subsampling of all transcript counts (cyan). Despite no significant changes in the weighted $\log_2(\mathrm{FC})$ and the corresponding weighted *t*-test values (shown on top), the violin plots look very different.

**Average and variance calculation in multiple sample experiments**

In multiple sample experiments, additional averaging over multiple samples must be performed

$$\bar{n}_Z = \sum_{s=1}^{K} \bar{n}_{Z,s} W_{Z,s} \ , \qquad (S19)$$

where

$$\bar{n}_{Z,s} = \sum_{i=1}^{N_s} n_{Z,i,s} w_{Z,i,s} \ , \qquad (S20)$$

$n_{Z,i,s}$ and $w_{Z,i,s}$ are $n_{Z,i}$ and $w_{Z,i}$ values for samples $s = 1,2\ldots K$, $N_s$ is the number of cells in sample $s$ and

$$W_{Z,s} = \frac{1}{Var(\bar{n}_{Z,s})} \Big/ \sum_{s=1}^{K} \frac{1}{Var(\bar{n}_{Z,s})} \qquad (S21)$$

is the statistical weight of sample $s$ ($\sum_s W_{Z,s} = 1$). Ignoring differences in statistical weights of different samples may be just as problematic as neglecting variations in statistical weights of individual cells.

Eqs. (*S*19),(*S*21) are identical to Eqs.(*S*2),(*S*3), yet here $Var(\bar{n}_{Z,s})$ is given by

$$Var(\bar{n}_{Z,s}) = \langle(\bar{n}_{Z,s} - \langle\bar{n}_{Z,s}\rangle)^2\rangle + \mathcal{V}_Z = \frac{1}{1 - \sum_{i=1}^{N_s} w_{Z,i,s}^2} \sum_{i=1}^{N_s} w_{Z,i,s}^2 (n_{Z,i,s} - \bar{n}_{Z,s})^2 + \mathcal{V}_Z \qquad (S22)$$

where $\mathcal{V}_Z$ is an analogue of $V_Z$ in Eq. (*S*5) and we used Eq. (*S*10) for $\langle(\bar{n}_{Z,s} - \langle\bar{n}_{Z,s}\rangle)^2\rangle$. Using the same logic as for Eq. (*S*17), we arrive at

$$\sum_{s=1}^{K} \frac{1 - \sum_{i=1}^{N_s} w_{Z,i,s}^2}{\sum_{i=1}^{N_s} w_{Z,i,s}^2 (n_{Z,i,s} - \bar{n}_{Z,s})^2 + \left(1 - \sum_{i=1}^{N_s} w_{Z,i,s}^2\right)\mathcal{V}_Z} = \frac{1 - \sum_{s=1}^{K} W_{Z,s}^2}{\sum_{s=1}^{K} W_{Z,s}^2 (\bar{n}_{Z,s} - \bar{n}_Z)^2}, \qquad (S23)$$

where $n_{Z,i,s}$ are measured, $w_{Z,i,s}$ are calculated by numerically solving Eq. (*S*17) or determined from ANOVA ICC for each individual sample *s*, $\bar{n}_{Z,s}$ are calculated from Eq. (*S*20), and $\mathcal{V}_Z$ and $W_{Z,s}$ are calculated by numerically solving Eqs. (*S*21)-(*S*23). The values of $W_{Z,s}$ can then be used for determining $\log_2(\mathrm{FC})$ and the corresponding *p*-values from the weighted *t*-test. R functions implementing all these calculations for Seurat objects are available for download from https://github.com/sergeyleikin/sc-sp-RNASeq. Testing of the robustness of this approach and its comparison with previously described pseudo-bulk calculations are described in the main text of the paper.

**Chi-squared test and its combination with weighted *t*-test**

The weighted *t*-test with statistical weights calculated from Eq. (*S*17) and Eqs. (*S*21)-(*S*23) for multiple sample experiments is the minimal assumptions approach to evaluating the significance of scRNASeq results. It is more robust and general than other approaches, yet it is not perfect either. Like previous scRNASeq data analysis models and models of cluster-randomized experiments, it still relies on individual cells being random technical replicates and individual samples being random biological replicates. The weighted *t*-test and all these models *conceptually* assume that the sequenced cells are randomly selected representatives of larger cell populations. In scRNASeq experiments, this is only partially true. Even in most careful experiments, at least a small fraction of cells is expected to be dramatically altered from their original state by the cell isolation process, e.g., resulting in apoptosis initiation. Expression of some genes in these cells may be orders of magnitude different from all other cells in the population and not random. As a result, even the most robust weighted *t*-test may detect these genes as differentially expressed, which is not desirable.

To mitigate this problem, we suggest using a $\chi^2$ test for the contingency Table 1 together with the weighted *t*-test.

| | **Cells A** | **Cells B** |
|---|---|---|
| **Gene *Z*** | $N_{Z,A}$ | $N_{Z,B}$ |
| **All other genes** | $N_A - N_{Z,A}$ | $N_B - N_{Z,B}$ |

**Table 1. Contingency table for $\chi^2$ test in scRNASeq.** $N_{Z,Y} = \sum_{i \in Y} N_{Z,i}$, $N_Y = \sum_Z N_{Z,Y}$, $Y = A, B$.

The benefit of this approach is that the $\chi^2$ test does not assume individual cells to be properly randomized replicates. Instead, it calculates the *p*-values only for the aggregated counts in specific cells sequenced in the scRNASeq experiment without generalizing it to all cells of the same type in the sample (see the derivation below). However, $n_{Z,i}$ variance within the sequenced cells may be smaller than the variance in the overall population of cells of the same type (even within the same sample). Therefore, positive findings of the $\chi^2$ test may not mean differential expression in the latter population. At the same time, negative findings of the $\chi^2$ test mean low likelihood of differential expression in the sequenced cells, helping to eliminate weighted *t*-test positives related to sampling bias. Therefore, performing the $\chi^2$ test and weighted *t*-test for all genes may be beneficial for reducing type I errors caused by experimental cell selection and processing bias.

Briefly, the validity of the $\chi^2$ test for this specific application can be proven as follows. This test for the contingency table in Table 1 is based on a $\chi^2$ distribution for

$$X^2 = \sum_{m=1}^{2}\sum_{n=1}^{2}\frac{\left(O_{m,n} - E_{m,n}\right)^2}{E_{m,n}} . \qquad \text{(S24)}$$

Here $O_{m,n}$ are the observed values in the contingency table and $E_{m,n}$ are the expected values of $O_{m,n}$ at $N_{Z,A}/N_A = N_{Z,B}/N_B$ ($\log_2(\mathrm{FC}) = 0$). $X^2$ has the $\chi^2$ distribution in the asymptotic limit of large $O_{m,n}$ when $Var(O_{m,n}) = E_{m,n}$ [18]. This simply requires $N_{Z,A}$ or $N_{Z,B}$ to be sufficiently large (our preference is $> 30 - 50$). Provided that this condition is satisfied, the accuracy of the $\chi^2$ test hinges only on how well $Var(N_Z)$ is approximated by $\langle N_Z \rangle$. At $Var(N_Z) < \langle N_Z \rangle$, the test becomes conservative. At $Var(N_Z) > \langle N_Z \rangle$, it becomes anti-conservative.

Taking into account that $N_{Z,i} = N_i n_{Z,i}$ and therefore $Var(N_{Z,i}) = N_i^2 \langle \left(n_{Z,i} - \langle n_{Z,i} \rangle\right)^2 \rangle$, similar to the derivation of Eq. (*S*6) we find that

$$Var(N_{Z,i}) = N_i \langle \langle n_{Z,i} \rangle_i \left(1 - \langle n_{Z,i} \rangle_i\right) \rangle \cong \langle N_{Z,i} \rangle \qquad \text{(S25)}$$

as long as sequencing of each transcript *j* in the cell *i* can be considered an independent transcript sampling event. Note that here the averaging indicated by the symbol $\langle\ \rangle$ is performed only over statistical trajectories that begin after encapsulation of each cell *i* into a droplet for single cell sequencing and does not include averaging over possible cell sampling trajectories. Therefore, $Var(N_{Z,i})$ does not include the variance associated with cell sampling, which is described by $V_Z$ in Eq. (*S*6).

Because each cell *i* is encapsulated in an independent droplet, the sequencing of its transcripts is independent from sequencing of transcripts in other cells. Therefore,

$$Var(N_Z) = \sum_i Var(N_{Z,i}) \cong \sum_i \langle N_{Z,i} \rangle = \langle N_Z \rangle . \qquad \text{(S26)}$$

This proves that the $\chi^2$ test provides an accurate estimate for the probability of the observed $N_{Z,A}$ and $N_{Z,B}$ at $N_{Z,A}/N_A = N_{Z,B}/N_B$. However, this is the probability only for the cells sequenced by NGS. It bears reminding that $Var(N_Z)$ for statistical trajectories that begin at the sample stage before cell isolation and encapsulation into droplets may be larger, even significantly larger. Thus, the $\chi^2$ test should not be used for inferring the *p*-value for a larger population of cells (even from the same sample).

**Sampling and technical noise in spRNASeq**

Aside from spatial information (Fig. 6), the output of sequencing based spRNASeq assays is like scRNASeq, i.e. $N_{Z,i}$ matrix of gene $Z$ counts in a spatial bin $i$. In contrast to scRNASeq, however, a single bin $i$ is likely to include transcripts from multiple cells, and adjacent bins may contain transcripts from the same cell. As a result, physics of transcript sampling and technical noise in spRNASeq depends on the spatial resolution of the assay.

At low resolution (bin size ~ 50 μm or larger), adjacent bins can be considered independent. DGE analysis in such assays is similar to scRNASeq. Provided that enough bins are included in the analysis (preferably ~ 100 or more), the approach described in the main text can be utilized without modifications. The only difference is that the bins must be interpreted as variable composition mixtures of cells. Note that relative count normalization may not be the best choice for some applications of these assays. For instance, normalization based on cell-type-specific endogenous control genes may enable analysis of gene expression in specific cell types despite the lack of single cell resolution. Approaches to such normalization are, however, context specific and are beyond the scope of the present paper.

At high spatial resolution (bin size ~ 10 μm or smaller), transcript counts in adjacent bins are likely to be correlated, precluding treatment of individual bins as independent replicates. One solution is correlation-based models, which are justifiable for isotropic tissues with similar correlations between bins in all spatial directions. Since we are more interested in anisotropic tissues like those shown in Fig. 6., we are not pursuing this approach.

Another solution is clustering of bins based on spatial localization (Fig. 6) or marker gene expression followed by analysis of aggregated transcript counts $\bar{n}_{Z,c} = \sum_{i \in c} N_{Z,i} / \sum_{i \in c} N_i$. Here $N_i = \sum_Z N_{Z,i}$ is the total number of counts in bin $i$. Transcript counts in sufficiently large clusters can be considered independent. Within such a cluster, the statistical weight of each bin is $w_{Z,i} = N_i / \sum_i N_i$, which is a good approximation at $\rho_Z(N_i - 1) \ll 1$ (see Eq. ($S8$)). Fig. 1B shows that this approximation works reasonably well for many genes even at $N_i$ between $10^4$ and $10^5$ in scRNASeq. We can then expect it to work even better in high resolution spRNASeq, where $N_i$ ~ 1000 or lower. At the current state of the technology, transcript counts in spRNASeq are simply not sufficient for analyzing biological variability within individual cell clusters.

DGE between two cell clusters can then be analyzed based on combining the weighted $t$-test and $\chi^2$ test as described in the main text. A more accurate DGE comparison can be made between two groups of multiple clusters, using weighted $t$-test like in multiple sample scRNASeq experiments (replacing $\bar{n}_{Z,s}$ and $W_{Z,s}$ in Eqs. ($S21$)-($S23$) with $\bar{n}_{Z,c}$ and the corresponding statistical weight $W_{Z,c}$ for each cluster). The same procedure can be further repeated for multiple samples.

**Computational benchmarking**

The goal of this paper is to demonstrate major artifacts caused by unneeded assumptions in commonly used scRNASeq and spRNASeq data analysis methods and show that a minimal-assumptions approach akin to that in cluster-randomized clinical trials largely eliminates such artifacts. To facilitate its understanding and testing, we provide the source code for functions implementing this minimal-assumptions approach in R programming language for Seurat data objects. The code is designed to ease understanding of the functions rather than to increase their computation speed. For instance, it performs all calculations sequentially in a loop for individual genes instead of vectorizing them. Much faster vectorized calculations could be easily accomplished for ANOVA-ICC-based analysis, yet this would be more challenging for iterative-ICC-based analysis. Thus, it makes more sense to first test multiple datasets for whether both ICC estimators produce similar results (as we suspect) or whether one of them performs better (which is possible). Such testing is beyond the scope of the present paper.

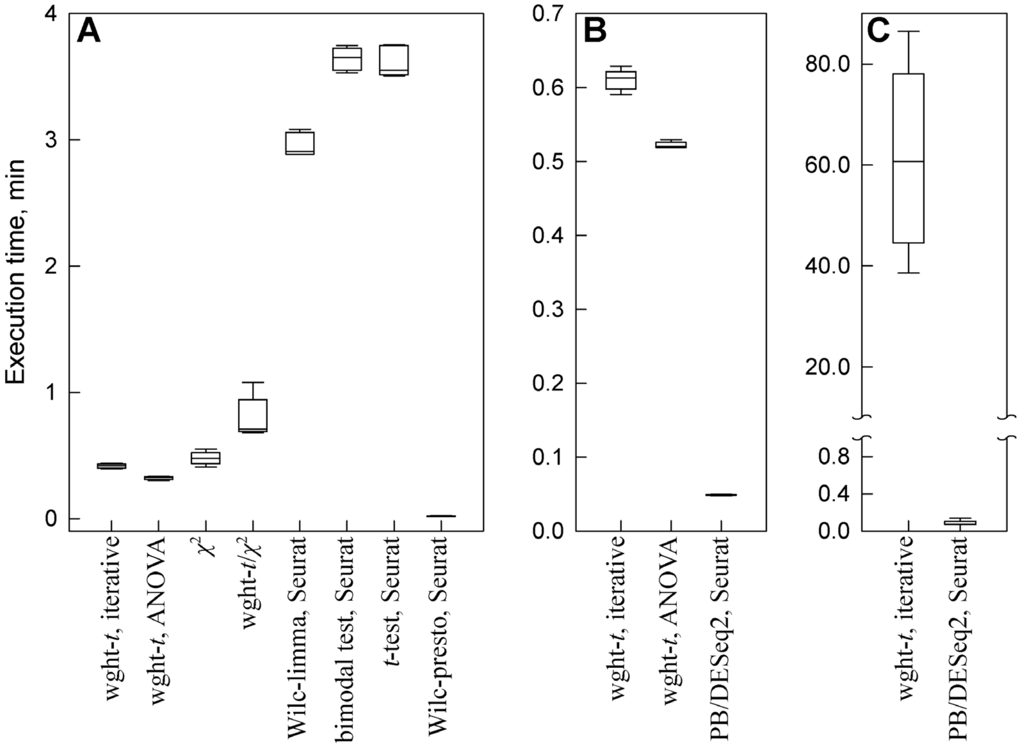


**Figure S1.6. Benchmarking DGE analysis functions. A.** Pairwise DGE analysis for 12,087 genes in 1096 glutamatergic neurons with 1,110 GABAergic neurons using weighted *t*-test with iterative ICC (wght-*t*, iterative) or ANOVA ICC (wght-*t*, ANOVA), $\chi^2$ test ($\chi^2$), weighted-$t/\chi^2$ test combination with iterative ICC (wght-$t/\chi^2$), limma implementation of the Wilcoxon test in Seurat (Wilc-limma, Seurat), bimodal test in Seurat (bimodal, Seurat), unweighted *t*-test in Seurat (*t*-test, Seurat), and presto implementation of the Wilcoxon test in Seurat (Wilc-presto, Seurat). **B.** Multi-sample DGE analysis (3 biological replicates) for 21,151 genes in 1,222 glutamatergic neurons with 1,303 GABAergic neurons using weighted *t*-test with iterative ICC, weighted *t*-test with ANOVA ICC, and DESeq2-based pseudobulk analysis implementation in Seurat (PB/DESeq2, Seurat). **C.** Multi-sample DGE analysis for 19,059 genes in 376,376 cells (total) from 3 wild type mice and 3 mutant mice. The data were collected using the Visium HD spRNASeq assay. For the purpose of this benchmarking, each spatial bin was treated as an individual cell. The weighted mean values and variances for each gene in each sample were calculated based on weighted averaging with ICC=0 (the same calculation speed as with ANOVA ICC) followed by weighted *t*-test with iterative ICC for the comparison of wild type and mutant samples. These samples are from a separate ongoing study and are used here only for benchmarking purposes. Since the mutation, tissue and cell identities are irrelevant for the benchmarking, they are not reported and have been removed from the shared dataset. **A,B,C.** The benchmarking was performed using a Dell Precision 7680 laptop workstation with 64 GB RAM, Intel i9-13950HX CPU, and NVIDEA RTX 3500 laptop GPU. The calculations were noticeably slowed down by mandatory IT security software running on the background, which appeared to be largely responsible for the observed execution time variations. The total execution time was measured in 5 simulations for each benchmark using the system.time() function in R. The box plots show the median, 10th, 25th, 75th, and 90th percentiles.

Nonetheless, we found the functions we provide to be fast enough for practical analysis of any datasets on a standard Dell Precision 7680 laptop workstation, despite the heavy burden of the IT security software running on the background (Fig. S1.6). In our hands, the DGE calculations by these functions were never even close to being the bottleneck of the overall data analysis. Much more time was consumed by standard preprocessing of raw sequencing data, data quality analysis, and assembly and annotation of Seurat objects. Typical analysis of DGE between two groups of several hundred cells was completed within a few seconds. More challenging DGE analysis of ~12,000 genes between ~1,100 glutamatergic neurons and ~1,100 GABAergic neurons was completed in less than 1 min (Fig. 1.6A). DGE analysis of ~20,000 genes for such neurons in 3 different datasets (~2,500 total neurons) took ~30 s (Fig. 1.6B). Even the most challenging analysis of DGE between two sets of data containing 3 samples each, ~ 20,000 genes, and ~ 400,000 cells took ~ 1 h (Fig. 1.6C). For comparison, the time to prepare for the latter analysis after the tissue samples were collected was ~ 3 weeks for sample processing and sequencing assay plus ~ 1 week for data processing to assemble the quality-assured, annotated Seurat object.

For completeness, we nevertheless compared the performance of our functions to the common Seurat workflow. When comparing just two groups of cells, our functions were faster than most tests built into the Seurat's FindMarkers function (Fig. S1.6A), including the common limma implementation of the Wilcoxon test (https://rdrr.io/bioc/limma/man/rankSumTestwithCorrelation.html). The only exception was the fast presto implementation of the Wilcoxon test (https://github.com/immunogenomics/presto), although the latter provided different results from the limma implementation. When comparing ~ 2,500 cells in multiple samples, our functions were noticeably slower than the DESeq2-based pseudobulk analysis in Seurat. The difference in performance became much more dramatic when multiple sample analysis was scaled to 400,000 cells (Fig. 1.6C), because the Seurat pseudobulk analysis was fully vectorized while our algorithm was not (as discussed above). In addition, the pseudobulk analysis simply aggregates counts while we fully analyze each sample, calculating ICC, mean, and variance for all genes.

Overall, we found that the nonoptimized implementation of our minimal-assumptions approach should be fast enough for practical analysis of any existing datasets. Much faster implementation based on vectorizing some of the calculations within our functions is possible, yet the ever-increasing computing power is likely to be sufficient for the ever-increasing data sizes even without it.

#### Appendix: Additional notes on derivation of Eq. (*S*6)

***Note 1:*** To address possible transcript sampling and identification failures in scRNASeq experiments, consider a more general form of Eq. ($S1$)

$$n_{Z,i} = \frac{1}{N_i}\sum_{j=1}^{N_i^{tot}} X_{Z,i,j} \qquad (S1')$$

where $N_i^{tot}$ is the unknown total number of transcripts in the cell $i$, including those identified and not identified by scRNASeq. $X_{Z,i,j} = 1$ for transcripts $j$ in the cell $i$ that are identified by scRNASeq as gene $Z$ mRNAs. $X_{Z,i,j} = 0$ for all other transcripts (successfully identified by scRNASeq as transcripts of other genes, sampled but not identified by failed sequencing, and not sampled at all). Since $X_{Z,i,j} \neq 0$ only for identified transcripts, we can limit the summation in Eq. ($S1'$) to the latter, i.e.,

$$\frac{1}{N_i}\sum_{j=1}^{N_i^{tot}} X_{Z,i,j} = \frac{1}{N_i}\sum_{j=1}^{N_i} X_{Z,i,j} \ , \qquad (S2')$$

recovering the original Eq. ($S$1).

***Note 2:*** Substitution of Eq. ($S$1) into Eq. ($S$5) yields

$$Var(n_{Z,i}) = \langle\left(\frac{1}{N_i}\sum_{j=1}^{N_i} X_{Z,i,j} - \langle\frac{1}{N_i}\sum_{j=1}^{N_i} X_{Z,i,j}\rangle_i\right)^2\rangle + V_Z = \langle\left(\frac{1}{N_i}\sum_{j=1}^{N_i}\left(X_{Z,i,j} - \langle X_{Z,i,j}\rangle_i\right)\right)^2\rangle + V_Z. \qquad (S3')$$

Mathematically, our assumption of random and independent transcript identification trials means that $\langle\left(X_{Z,i,j} - \langle X_{Z,i,j}\rangle_i\right)\left(X_{Z,i,k} - \langle X_{Z,i,k}\rangle_i\right)\rangle_i = 0$ at $j \neq k$. Therefore,

$$Var(n_{Z,i}) = \langle\frac{1}{N_i^2}\sum_{j=1}^{N_i}\left(X_{Z,i,j} - \langle X_{Z,i,j}\rangle_i\right)^2\rangle + V_Z. \qquad (S4')$$

After rewriting Eq. ($S$4') as

$$Var(n_{Z,i}) = \langle\frac{1}{N_i^2}\sum_{j\in\{X_{Z,i,j}=0\}}\left(X_{Z,i,j} - \langle X_{Z,i,j}\rangle_i\right)^2 + \frac{1}{N_i^2}\sum_{j\in\{X_{Z,i,j}=1\}}\left(X_{Z,i,j} - \langle X_{Z,i,j}\rangle_i\right)^2\rangle + V_Z, \qquad (S5')$$

where $\{X_{Z,i,j} = 0\}$ and $\{X_{Z,i,j} = 1\}$ are the sets of $j$ values with the corresponding $X_{Z,i,j}$, and using that

$$\langle n_{Z,i}\rangle_i = \frac{1}{N_i}\sum_{j=1}^{N_i}\langle X_{Z,i,j}\rangle_i = \langle X_{Z,i,j}\rangle_i \ , \qquad (S6')$$

we find

$$Var(n_{Z,i}) = \langle\frac{1}{N_i^2}\sum_{j\in\{X_{Z,i,j}=0\}}\langle n_{Z,i}\rangle_i^{\,2} + \frac{1}{N_i^2}\sum_{j\in\{X_{Z,i,j}=1\}}\left(1 - \langle n_{Z,i}\rangle_i\right)^2\rangle + V_Z, \qquad (S7')$$

Then, after substitution of

$$\sum_{j\in\{X_{Z,i,j}=1\}}^{N_i} 1 = N_i n_{Z,i} \quad , \qquad \sum_{j\in\{X_{Z,i,j}=0\}}^{N_i} 1 = N_i\left(1 - n_{Z,i}\right) \qquad (S8')$$

into Eq. ($S$7'), we arrive at

$$Var(n_{Z,i}) = \langle\frac{1}{N_i^2}\langle n_{Z,i}\rangle_i^{\,2} N_i\left(1 - n_{Z,i}\right) + \frac{1}{N_i^2}\left(1 - \langle n_{Z,i}\rangle_i\right)^2 N_i n_{Z,i}\rangle + V_Z. \qquad (S9')$$

Since the overall averaging can be performed in two steps, first averaging within each cell *i* and then across different cells, we can replace $n_{Z,i}$ in Eq. (*S*9*'*) with $\langle n_{Z,i}\rangle_i$ and simplify this equation to

$$Var(n_{Z,i}) = \langle\frac{\langle n_{Z,i}\rangle_i(1-\langle n_{Z,i}\rangle_i)}{N_i}\rangle + V_Z. \qquad (S10')$$

Next, using that $\langle\langle n_{Z,i}\rangle_i\rangle = \langle n_{Z,i}\rangle$ and

$$V_Z = Var(\langle n_{Z,i}\rangle_i) = \langle\langle n_{Z,i}\rangle_i{}^2\rangle - \langle\langle n_{Z,i}\rangle_i\rangle^2 \qquad (S11')$$

we arrive at

$$Var(n_{Z,i}) = \frac{\langle n_{Z,i}\rangle}{N_i} - \frac{Var(\langle n_{Z,i}\rangle) + \langle n_{Z,i}\rangle^2}{N_i} + V_Z = \frac{\langle n_{Z,i}\rangle}{N_i} - \frac{V_Z + \langle n_{Z,i}\rangle^2}{N_i} + V_Z \qquad (S12')$$

Finally, after taking into account that

$$\bar{n}_Z = \sum_{i=1}^{N} n_{Z,i}w_{Z,i} = \langle n_{Z,i}\rangle\ , \qquad (S13')$$

we arrive at Eq. (*S*6).

In other words, Eqs. (*S*6),(*S*7) are valid for any distributions of $X_{Z,i,j}$ in the asymptotic limit of large *N*, as long as the sampling of transcripts is random and independent. These distributions include but are not limited to the popular binomial, negative binomial, and zero-inflated negative binomial distributions. Because these equations are based on unknown $\bar{n}_Z$ and $V_Z$ rather than a defined probability distribution for $X_{Z,i,j} = 1$, they do not require assuming anything about cell transcripts that have not been identified by scRNASeq sampling. The unknown $\bar{n}_Z$ and $V_Z$ are then calculated directly from the experimental data without any additional assumptions by combining Eqs. (*S*2), (*S*8), and (*S*17), as described in the main text of this Supplement.

***<u>Note 3:</u>*** To illustrate how Eqs. $(S12')$ and $(S6)$ can be derived from common counting models, consider a textbook model of colored balls in baskets. Specifically, each cell *i* corresponds to a basket *i*, $N_{Z,i}$ corresponds to the number of balls with color *Z* drawn from that basket, $N_i$ corresponds to the total number of random draws from the basket *i*. The total numbers of color *Z* balls ($N_{Z,i}^{tot}$) and all balls ($N_i^{tot}$) in the basket *i* are not known, just like the total numbers of gene *Z* and all mRNA transcripts. The goal is to estimate

$$\bar{n}_Z = \langle N_{Z,i}^{tot}\ /N_i^{tot}\rangle \qquad (S14')$$

and $Var(N_{Z,i}^{tot}\ /N_i^{tot})$ based on the known $N_{Z,i}$ and $N_i$, so that the difference in the prevalence of color *Z* balls between two different sets of baskets can be evaluated based on the weighted *t*-test.

This model is described by the hypergeometric probability distribution for $N_{Z,i}$ that has the within-basket mean $\langle N_{Z,i}\rangle_i = N_i(N_{Z,i}^{tot}/N_i^{tot})$ and the variance

$$Var_i(N_{Z,i}) = N_i\left(\frac{N_{Z,i}^{tot}}{N_i^{tot}}\right)\left(\frac{N_i^{tot} - N_{Z,i}^{tot}}{N_i^{tot}}\right)\left(\frac{N_i^{tot} - N_i}{N_i^{tot} - 1}\right), \qquad (S15')$$

where $Var_i(N_{Z,i})$ is the variance within a single basket $i$. Then, using the law of total variance, we find that the total valiance of $n_{Z,i} = N_{Z,i}/N_i$ for all baskets with $N_i$ draws is

$$Var(n_{Z,i}) = \frac{\langle Var_i(N_{Z,i})\rangle}{N_i^2} + V_Z\,, \qquad (S16')$$

where $V_Z = Var(N_{Z,i}^{tot}/N_i^{tot})$ is unknown. After substituting $n_{Z,i}^{tot} = N_{Z,i}^{tot}/N_i^{tot}$ into Eq. ($S16'$), we find

$$Var(n_{Z,i}) = \frac{1}{N_i}\langle n_{Z,i}^{tot}(1-n_{Z,i}^{tot})(1-SR_i)\rangle + V_z\,, \qquad (S17')$$

where $SR_i = (N_i-1)/(N_i^{tot}-1)$ is the sampling ratio in the basket $i$. Taking into account that

$$\langle n_{Z,i}^{tot}(1-n_{Z,i}^{tot})\rangle = \langle n_{Z,i}^{tot}\rangle - \langle (n_{Z,i}^{tot})^2\rangle = \langle n_{Z,i}^{tot}\rangle - \langle n_{Z,i}^{tot}\rangle^2 - V_z\,, \qquad (S18')$$

we find

$$Var(n_{Z,i}) = \frac{\bar{n}_Z(1-\bar{n}_Z) + V_Z(N_i-1)}{N_i} - \frac{\langle SR_i n_{Z,i}^{tot}(1-n_{Z,i}^{tot})\rangle}{N_i}. \qquad (S19')$$

At $SR_i \ll 1$, i.e., when the sampling is random and independent, the last term in Eq. ($S19'$) is negligible and we recover Eqs. ($S12'$) and ($S6$). Then, at sufficiently large number of baskets, Eqs. ($S2$), ($S8$), and ($S17$) can be used to find the unknown values of $\bar{n}_Z$ and $V_Z$ without any additional assumptions based on matching the predicted and observed values of weighted average and variance.

The instructive value of this model is that it illustrates when and why one can analyze relative prevalence $n_{Z,i}$ and accurately estimate its average $\bar{n}_Z$ and variance $V_Z$ from the known $N_{Z,i}$ and $N_i$, even though $N_{Z,i}^{tot}$ and $N_i^{tot}$ are not known. In the context of this specific model, the key condition is the low sampling ratio ($N_i \ll N_i^{tot}$), which ensures random and independent sampling. At $N_i \sim N_i^{tot}$, the problem becomes more complex and may require additional assumptions. In the context of scRNASeq, $N_i \ll N_i^{tot}$ in all existing and foreseeable assays, so that such assumptions (which may not be justified) are not necessary. More generally, random and independent sampling is all that is needed to accurately estimate relative prevalence of a trait in a large group of independent population clusters (baskets in this example or cells in scRNASeq).

***Note 4:*** In a canonical sequence of Bernoulli trials, the positive outcome probability (rate) $p$ is the same in all trials, yielding binomial distribution. In the limit of infinite $N_i^{tot}$, Eq. ($S6$) can be derived by a straightforward application of the law of total variance to an ensemble of cells with fixed $N_i$ binomial trials each, where each cell has a different, but fixed $p$. However, Eq. ($S6$) is valid also when $p$ is variable rather than constant even within each cell (as our derivation in Note 2 above proves.) It is therefore instructive to illustrate this by using the binomial/Bernoulli distribution logic yet relaxing the requirement of constant within-cell $p$ to ensure more realistic description of scRNASeq.

Within a cell $i$, an individual trial $j$ that has the success probability $p_j$ is described by the Bernoulli distribution with $\langle X_{Z,i,j}\rangle_j = p_j$ and $Var_j(X_{Z,i,j}) = p_j(1-p_j)$. When $p_j$ varies for individual transcripts of the same gene $Z$ even within the same cell, the within-cell variance of $X_{Z,i,j}$ is

$$Var_i(X_{Z,i,j}) = \langle (X_{Z,i,j} - p_j + p_j - p)^2 \rangle_i = \langle (X_{Z,i,j} - p_j)^2 \rangle_i + \langle (p_j - p)^2 \rangle_i$$
$$= \langle Var_j(X_{Z,i,j}) \rangle_i + Var_i(p_j) = \langle p_j(1-p_j) \rangle_i + Var_i(p_j) = p(1-p)\,, \qquad (S20')$$

where $p = \langle p_j \rangle_i$ is the within-cell average rate, and we used that $\langle x \rangle_i = \langle \langle x \rangle_j \rangle_i$ and $\langle p_j^2 \rangle_i = \langle p_j \rangle_i^2 + Var_i(p_j)$.

For $N_i$ random and independent trials we can then calculate the across-cell variance of $n_{Z,i}$ as

$$Var(n_{Z,i}) = \frac{\langle Var_i(N_{Z,i}) \rangle}{N_i^2} + V_Z = \frac{\langle \sum_{j=1}^{N_i} Var_i(X_{Z,i,j}) \rangle}{N_i^2} + V_Z = \frac{\langle p(1-p) \rangle}{N_i} + V_Z\,. \qquad (S21')$$

Here $\langle\ \ \rangle$ indicates averaging across all cells (as before), $V_Z = Var(p)$, we used the law of total variance (Eq. ($S5$)), and we used that the variance of a sum of independent random variables is equal to the sum of variances of these variables. After substitution of $\langle p^2 \rangle = \langle p \rangle^2 + Var(p)$ into Eq. ($S21'$), we find

$$Var(n_{Z,i}) = \frac{\langle p \rangle (1 - \langle p \rangle) + V_Z(N_i - 1)}{N_i} \qquad (S22')$$

Eq. ($S22'$) has the same level of generality as our original Eq. ($S6$). Moreover, since $\langle p \rangle = \bar{n}_Z$, the two equations are actually identical. The key observations here are: (a) The only difference between Eq. ($S22'$) and the corresponding expression for binomially distributed $N_{Z,i}$ is that $\langle p \rangle$ averages $p$ variations both within and across the cells rather than just $p$ variations across the cells. (b) The variance for a single independent trial with a binary outcome and random probability $p$ is $\langle p \rangle(1 - \langle p \rangle)$, i.e., it depends only on $\langle p \rangle$ but not on the variability of $p$ (Eq. ($S20'$) and Eq. ($S22'$) with $N_i = 1$).

These observations suggest that Eq. ($S19'$) may work beyond the hypergeometric distribution not only when $SR_i \to 0$ but also at larger $SR_i$. However, this is a nontrivial question because at $SR_i \sim 1$ the expected variance is no longer a sum of variances for individual trials. Analysis of this question is beyond the scope of our study, since the case of $N_i \sim N_i^{tot}$ is not practically relevant for scRNASeq.

#### References


1. Amezquita, R.A., Lun, A.T.L., Becht, E., Carey, V.J., Carpp, L.N., Geistlinger, L., Marini, F., Rue-Albrecht, K., Risso, D., Soneson, C., et al. (2020). Orchestrating single-cell analysis with Bioconductor. Nat Methods 17, 137-145. 10.1038/s41592-019-0654-x.
2. Hafemeister, C., and Satija, R. (2019). Normalization and variance stabilization of single-cell RNA-seq data using regularized negative binomial regression. Genome Biol 20, 296. 10.1186/s13059-019-1874-1.
3. Lause, J., Berens, P., and Kobak, D. (2021). Analytic Pearson residuals for normalization of single-cell RNA-seq UMI data. Genome Biol 22, 258. 10.1186/s13059-021-02451-7.
4. Townes, F.W., Hicks, S.C., Aryee, M.J., and Irizarry, R.A. (2019). Feature selection and dimension reduction for single-cell RNA-Seq based on a multinomial model. Genome Biol 20, 295. 10.1186/s13059-019-1861-6.

5. Svensson, V. (2020). Droplet scRNA-seq is not zero-inflated. Nat Biotechnol 38, 147-150. 10.1038/s41587-019-0379-5.
6. Goodwin, S., McPherson, J.D., and McCombie, W.R. (2016). Coming of age: ten years of next-generation sequencing technologies. Nat Rev Genet 17, 333-351. 10.1038/nrg.2016.49.
7. Islam, S., Zeisel, A., Joost, S., La Manno, G., Zajac, P., Kasper, M., Lonnerberg, P., and Linnarsson, S. (2014). Quantitative single-cell RNA-seq with unique molecular identifiers. Nat Methods 11, 163-166. 10.1038/nmeth.2772.
8. Shapiro, E., Biezuner, T., and Linnarsson, S. (2013). Single-cell sequencing-based technologies will revolutionize whole-organism science. Nat Rev Genet 14, 618-630. 10.1038/nrg3542.
9. Fleiss, J.L. (1999). The Design and Analysis of Clinical Experiments (John Wiley & Sons).
10. Hartung, J., Knapp, G., and Sinha, B. (2008). Statistical Meta-Analysis with Applications (Wiley-Interscience).
11. Hayes, R.J., and Moulton, L.H. (2022). Cluster Randomized Trials (CRC Press).
12. Bevington, P.R., and Robinson, D.K. (2002). Data Reduction and Error Analysis for the Physical Sciences, Third Edition Edition (McGraw-Hill).
13. Feynman, R.P. (1998). Statistical Mechanics: A Set of Lectures (CRC Press).
14. Blitzstein, J.K., and Hwang, J. (2015). Introduction to Probability (CRC Press).
15. Cochran, W.G. (1939). The use of the analysis of variance in enumeration by sampling. J Am Stat Ass 34, 492-510.
16. Elston, R.C., Hill, W.G., and Smith, C. (1977). Query: estimating "heritability" of a dichotomous trait. Biometrics 33, 231-236.
17. Ridout, M.S., Demetrio, C.G., and Firth, D. (1999). Estimating intraclass correlation for binary data. Biometrics 55, 137-148. 10.1111/j.0006-341x.1999.00137.x.
18. Ramachandran, K.M., and Tsokos, C.P. (2021). Mathematical Statistics with Applications in R, Third Edition Edition (Academic Press).
