## Supplement 2 for "Differential expression analysis in single cell and spatial RNASeq without model assumptions"

### Additional supplemental figures

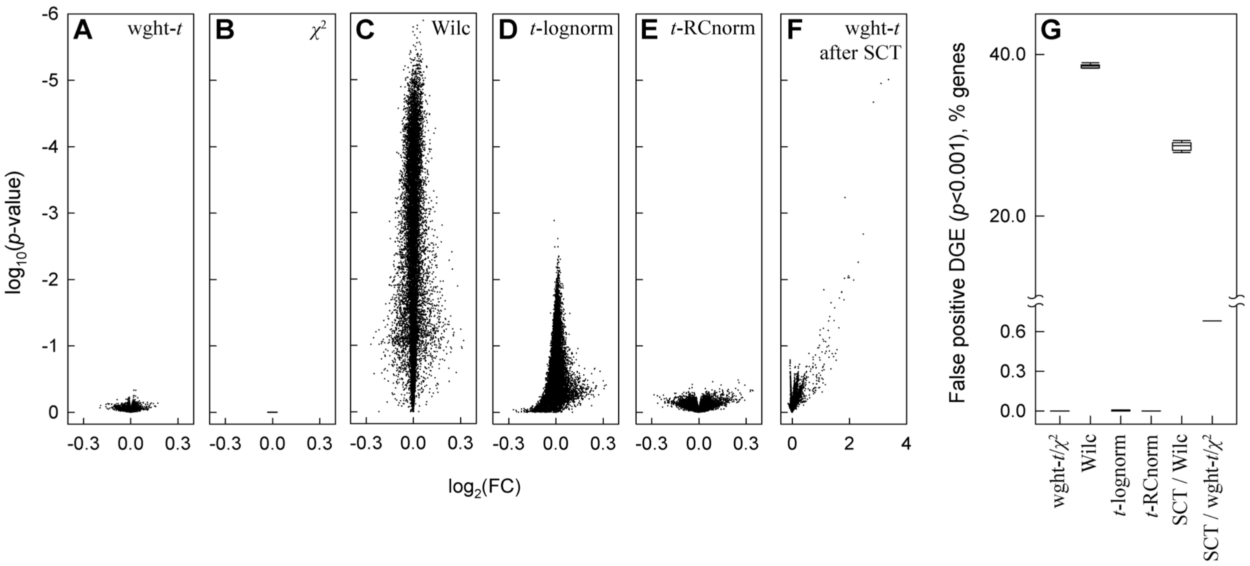


**Figure S2.1. False DGE discovery in glutamatergic neurons by common data analysis models after 60% UMI subsampling. A-E.** Volcano plots comparing original data with a single 60% UMI subsampling simulation using weighted *t*-test (**A**), $\chi^2$-test (**B**), Seurat's FindMarkers function default Wilcoxon test (**C**), Seurat's FindMarkers optional *t*-test for logarithmically normalized data (**D**), and *t*-test for RC normalized data (**E**). **F.** Volcano plot comparing SCTransform rescaled original data with SCTransform rescaled subsampled data using weighted *t*-test. **G.** Fraction (%) of 12,911 genes with more than 30 total transcript counts in 1096 glutamatergic neurons, which was falsely discovered by indicated tests as differentially expressed with $p < 0.001$ after 10 simulations. Note significant false DGE with very large $\log_2(\text{FC})$, which was introduced by SCTransform-based data rescaling (**F**).

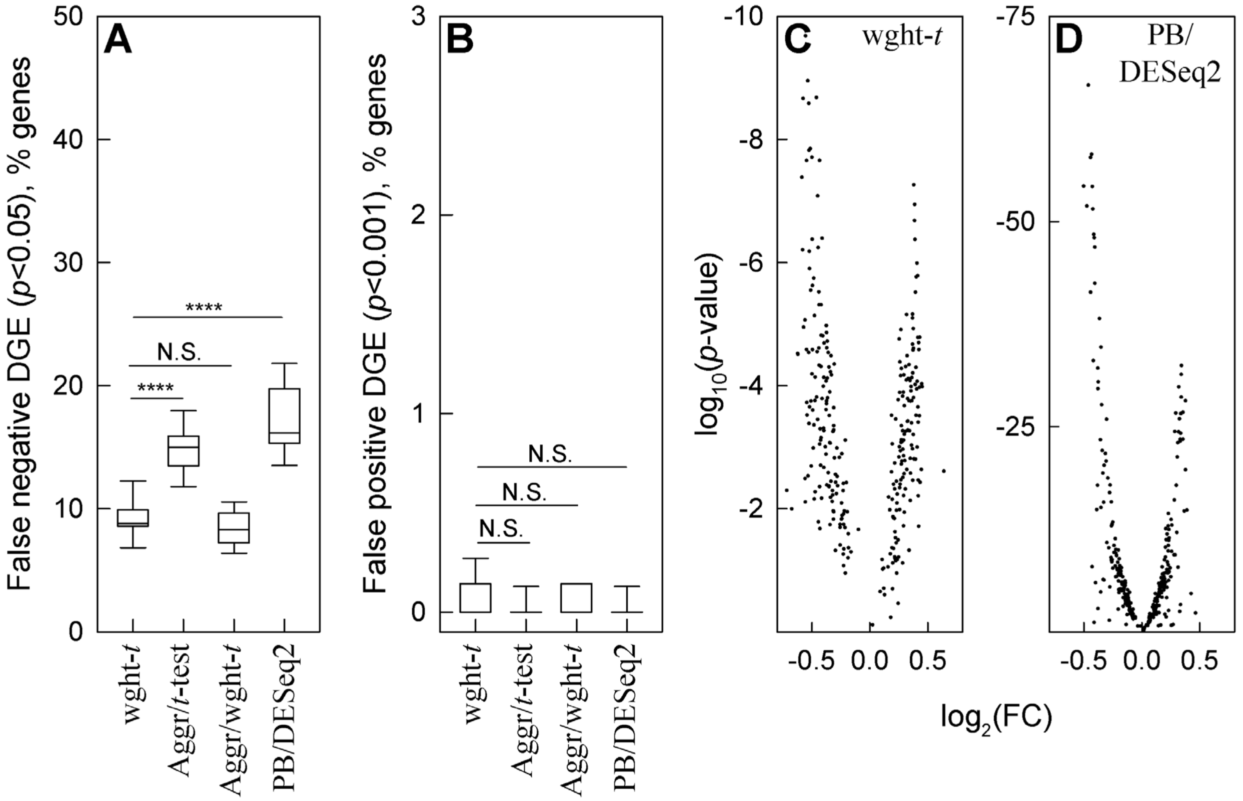


**Figure S2.2. Detection of DGE superimposed with random noise in multiple sample simulations.** Two 5-sample sets were generated as in Fig.5. Random noise and changes in expression were also generated as in Fig. 5, except 150 genes were modified by a 30% decrease in expression and 150 genes were modified by 30% increase in expression. **A.** False negative DGE among 300 genes with altered expression. **B.** False positive DGE among 700 genes modified only by noise. **C** and **D.** Volcano plots for 300 genes with altered expression analyzed by weighted *t*-test (C) and PB/DESeq2 (D). In panels A and B, **** means $p < 0.0001$, *** – $p < 0.001$, * – $p < 0.05$, N.S. – not significant.
